# Optimizing root phenotyping: Assessing the impact of camera calibration on 3D root reconstruction

**DOI:** 10.1101/2025.03.09.642238

**Authors:** Peter Pietrzyk, Suxing Liu, Alexander Bucksch

## Abstract

Accurate 3D reconstruction is essential for high-throughput plant phenotyping, particularly for studying complex structures such as root systems. While photogrammetry and Structure from Motion (SfM) techniques have become widely used for 3D root imaging, the camera settings used are often underreported in studies, and the impact of camera calibration on model accuracy remains largely underexplored in plant science. In this study, we systematically evaluate the effects of focus, aperture, exposure time, and gain settings on the quality of 3D root models made with a multi-camera scanning system. We show through a series of experiments that calibration significantly improves model quality, with focus misalignment and shallow depth of field (DoF) being the most important factors affecting reconstruction accuracy. Our results further show that proper calibration has a greater effect on reducing noise than filtering it during post-processing, emphasizing the importance of optimizing image acquisition rather than relying solely on computational corrections. This work improves the repeatability and accuracy of 3D root phenotyping by giving useful calibration guidelines. This leads to better trait quantification for use in crop research and plant breeding.

## 1. Background

Plant phenotyping has undergone a significant transformation thanks to advanced imaging and computer vision techniques (1, 2, 3, 4), particularly to exploit the information encoded in a plant’s morphology (5) from a whole plant level (6) to single-cell structures like unicellular root hairs (7). Within this area of research, high-resolution three-dimensional (3D) reconstruction of plants has emerged as a powerful tool for studying plant architecture (8, 9, 10), growth dynamics (11, 12), and responses to various environmental or agricultural stimuli (13). Among technologies to create such 3D models of plants are Lidar (14, 15), laser light-section (16), depth cameras (9), X-ray Computed Tomography (CT) (17, 18), as well as methods based on collections of two-dimensional (2D) images (19, 20). These non-destructive and high-throughput approaches provide valuable insights into crop yield and plant resistance to biotic and abiotic stress, whether in the field or indoor, spanning scales from plant organs to individual plants to whole plots (21).The insights include key information improving crop productivity and plant resilience to biotic and abiotic stress.

Image-based phenotyping of crop roots (22), in particular, has gained prominence due to the pivotal role of roots in nutrient uptake, water acquisition, and overall plant performance (23, 24, 25). The integration of root phenotyping with 3D measurement techniques can significantly enhance our ability to capture and analyze root traits and to identify corresponding genes (26). While X-ray CT has been used to study root traits of plants grown in pots, in recent years 3D reconstruction from 2D images has been increasingly utilized to measure traits of excavated roots from plants grown in the field. By utilizing camera rigs equipped with multiple high-resolution 2D color cameras and employing techniques from close-range photogrammetry, researchers can now generate detailed 3D models of plant roots in a cost-efficient way (27, 28, 29). However, the success of this phenotyping technique heavily depends on the quality of the generated 3D point clouds, as it forms the foundation for further processing steps such as skeletonization and trait extraction (30).

Central to the success of 3D reconstruction of plants, both roots and shoots, from 2D images is the careful calibration of the camera itself, i.e. intrinsic and extrinsic parameters, and the adjustment of camera settings, as for example focus, aperture, shutter speed, and gain. The two types of calibration play distinct but complementary roles in producing high-quality 3D points clouds for phenotyping.

Camera calibration refers to the estimation and correction of intrinsic parameters, which include focal length, principal point, and lens distortion, and extrinsic parameters that are related to the camera’s position and orientation in space (31, 32). This process ensures that the geometry of a scene is accurately defined and is often performed automatically using methods such as Structure from Motion (SfM) (33). Proper calibration is essential for ensuring that the 3D reconstruction reflects the true spatial relationship within the scene. The capability to automatically self-calibrate cameras using algorithms such as SfM, eliminating the need for manual camera calibration by the user, is an appealing feature for large-scale plant phenotyping and has allowed many researchers to use 3D reconstruction of plants at organ, individual plant, and plot scale.

On the other hand, camera settings—such as focus, aperture, shutter speed, and gain—do not directly alter the geometric calibration but play a crucial role in optimizing image quality (see Box 1). These settings affect the clarity, sharpness, and exposure of the captured images, which in turn influences the accuracy of the resulting 3D point cloud (34). Many of these settings can be interdependent. For example, changing either the root or the camera position will require readjustment of the camera’s focus. Similarly, changing the aperture (f-value) from a wide-open iris with f/2 to a small iris with f/8 will increase DoF and thus the overall sharpness of the imaged root. At the same time, the closed iris allows less light to reach the camera sensor and thus results in darker images, which must be compensated for either with longer exposure times, higher gain or additional light to obtain the same image brightness. As a result, it is crucial to consider the effect of each setting on image quality and define requirements to adjust these.

### Box 1: Definition of common camera settings

**Focus** addresses the distance between the camera’s sensor and the focal point, where the subject appears sharpest in the image. A camera focused on the root ensures that specific regions of the roots, especially fine root structures, are sharp and detailed. Poor focus can blur essential features, making it difficult to reconstruct accurate 3D models.

**Aperture** controls the amount of light entering the camera and affects the(DoF, i.e. the range of the scene that appears sharp. A small aperture (high f-number) increases the DoF, ensuring more of the plant is in focus, which is crucial for capturing detailed structures in complex root systems. However, a smaller aperture (small f-number) reduces light, requiring adjustments in other settings like shutter speed to maintain brightness, and can result in blur caused by diffraction.

**Shutter speed** determines the exposure time—the duration that the camera sensor is exposed to light. In dynamic environments, such as phenotyping platforms where cameras or plants may move, a fast shutter speed prevents motion blur. However, a faster shutter reduces the amount of light reaching the sensor, potentially darkening the image.

**Gain (ISO)** controls the camera sensor’s sensitivity to light. Increasing gain brightens the image in low-light conditions but introduces noise, which can obscure fine plant details. Lower gain settings are preferred to minimize noise and preserve image clarity for accurate 3D reconstructions.

Despite their critical role in image quality, camera settings are often underreported in plant phenotyping studies (28, 35). Some studies evaluate the effect of other variables, such as image resolution or viewing angle, on model quality but either do not report aperture, shutter speed and gain at all or use vague language (36, 37, 38). Suboptimal camera settings can result in deteriorated 3D reconstructions, making the subsequent skeletonization and root trait extraction less reliable. The insufficient reporting of these settings in turn can impact the reproducibility of the results, as has already been observed with the lack of documentation and transparency in the development of other imaging pipelines for plant phenotyping (39).

Considering these factors, this study aims to guide non-experts through the camera setup process and to illustrate the influence of specific camera settings on the quality and accuracy of 3D point clouds of roots. By using a multi-view 3D system with ten cameras rotating around an excavated maize root crown (27), our research aims to explore how variations in focus, aperture, shutter speed, gain and other camera settings affect the resulting 3D models. Given the dense and highly occluded nature of maize root crowns with fine structures in the order of 100 microns in diameter, achieving accurate reconstructions is challenging but essential for precise phenotyping. The pipeline for 3D reconstruction incorporates COLMAP to generate a 3D point cloud and a noise filtration process to reduce noise and eliminate blunders caused by the reconstruction. By assessing the impact of camera calibration on 3D point cloud quality, this study evaluates the robustness of 3D reconstruction across different camera set-ups and challenging working environments, ultimately contributing to more reliable and reproducible phenotyping pipelines in crop science.

## 2. Materials and Method

### 2.1 Root Samples

We selected 12 samples of contrasting maize (*Zea mays*) root architectures (Figure 1). The plants were grown in Hagerstown silt loam soil and harvested 81 d after planting at the Pennsylvania State University’s Russell E. Larson Agricultural Research Center in August 2018. After harvesting, we removed the shoots above all root-producing nodes and air-dried them on a greenhouse bench. Roots were transported to the University of Georgia in Athens, GA, and were then stored and imaged for this publication in February 2023.

**Figure 1:**
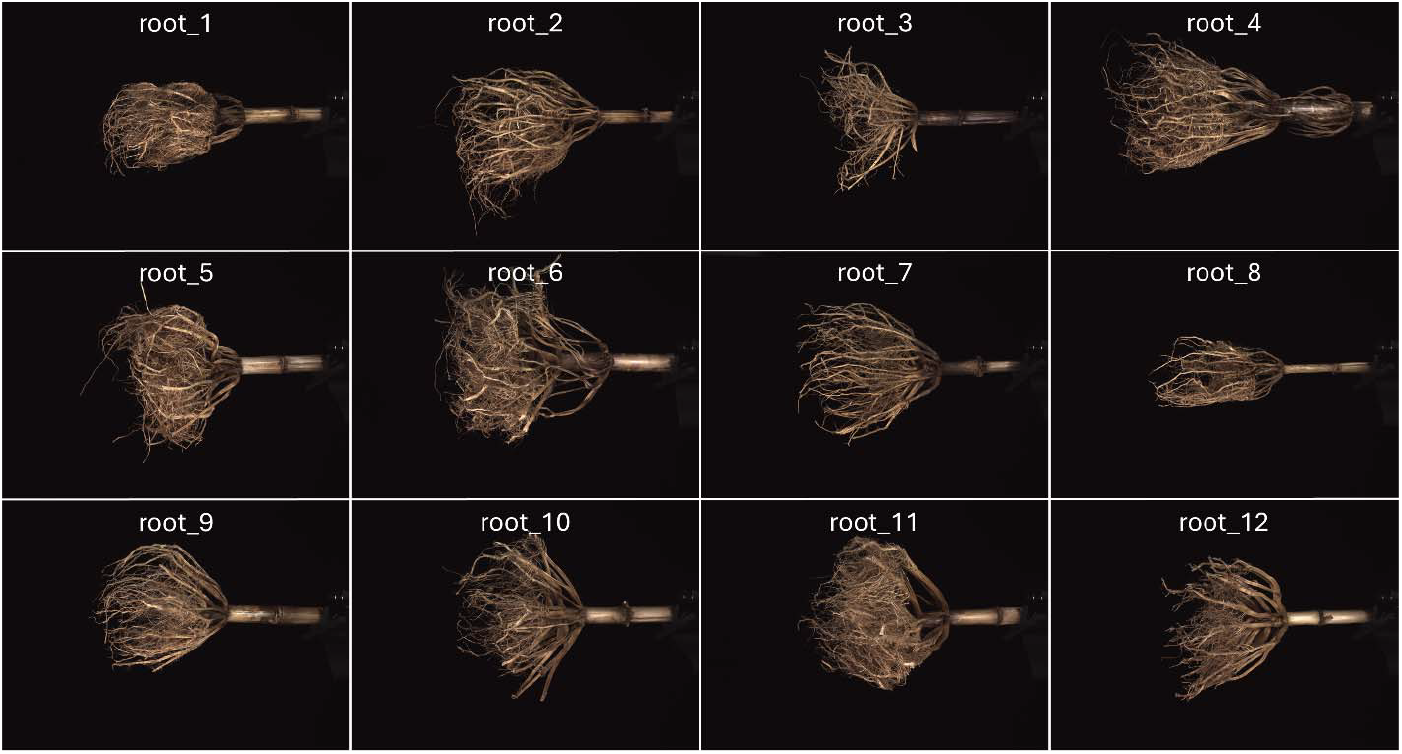
Overview of all 12 selected roots. All images were taken using the calibrated scanner setup and have the same scale.

### 2.2 3D Root Scanner

The 3D root scanner is described in (27) and consists of 10 cameras mounted on a round metal frame that rotates around the root sample. The scanner uses The Imaging Source DFK 27BUJ003 color cameras with 1/2.3 inch Aptina CMOS MT9J003 sensors. This camera has a maximum resolution of 10.7 megapixels, and its shutter speed ranges from ^1^/_20,000_ to ¼ sec, gain from 0 to 12 dB, and white balance from -2 to 6 dB. The cameras are further equipped with The Imaging Source TCL 0616 5 Mega Pixel lenses with 1/1.8 inch format, fixed 6 mm focal length, aperture range from f/1.6 to f/16, and a minimum object distance of 0.1 m.

### 2.3 Adjustment of Camera Settings

To increase 3D root model quality and accuracy of measured root traits, we developed a procedure to set up the scanner, which included the position and orientation of the root sample itself and settings for the cameras and environment as listed in Table 1. Other factors related to the computation of the 3D model, such as intrinsic and extrinsic parameter calibration with COLMAP and its meta-parameters, were not considered. During the calibration process, we used the software tcam-capture by The Imaging Source to interactively view and save images and to test camera settings of individual cameras. We turned off the automatic adjustment of exposure and gain during this process and manually set all other camera parameters.

**Table 1:** List of parameters that were adjusted during the set-up, together with an indication of whether parameters were calibrated and analyzed for their effect on model quality.

| Object | Setting | Calibrated and tested |
| --- | --- | --- |
| Root | Position | No |
|  | Orientation | No |
| Camera | Position | Yes |
|  | Orientation | Yes |
|  | Focus | Yes |
|  | Aperture | Yes |
|  | Exposure time | Yes |
|  | Gain | Yes |
|  | White balance | No |
|  | Image format/quality | Yes |
| Environment | Light source (position and power) | No |

### 2.4 Experiments

#### 2.4.1 Experiment 1

We performed two experiments to determine the effect of scanner parameters on the quality of point clouds. An overview of the parameters used in each experiment is given in Table 2. In Experiment 1, we performed two tests (E1#1 and E1#2), each with all twelve roots, resulting in a total of 24 scans. The first test uses a calibrated scanner setup (focused on center, aperture of f/4, exposure time of 1/30 sec and 0 dB gain), while for the second test we simulated an uncalibrated setup using the settings from Table 2. For the uncalibrated scan we used an aperture of f/2 to reduce DoF, set the focal point 10cm behind or in front of the scanner center alternating between cameras, and increased gain by 12 dB (equivalent to a 16-fold exposure) to introduce speckle. Due to the higher gain and the wider aperture, we had to reduce exposure time to 1/1000 sec, which still resulted in overexposed images.

**Table 2:**
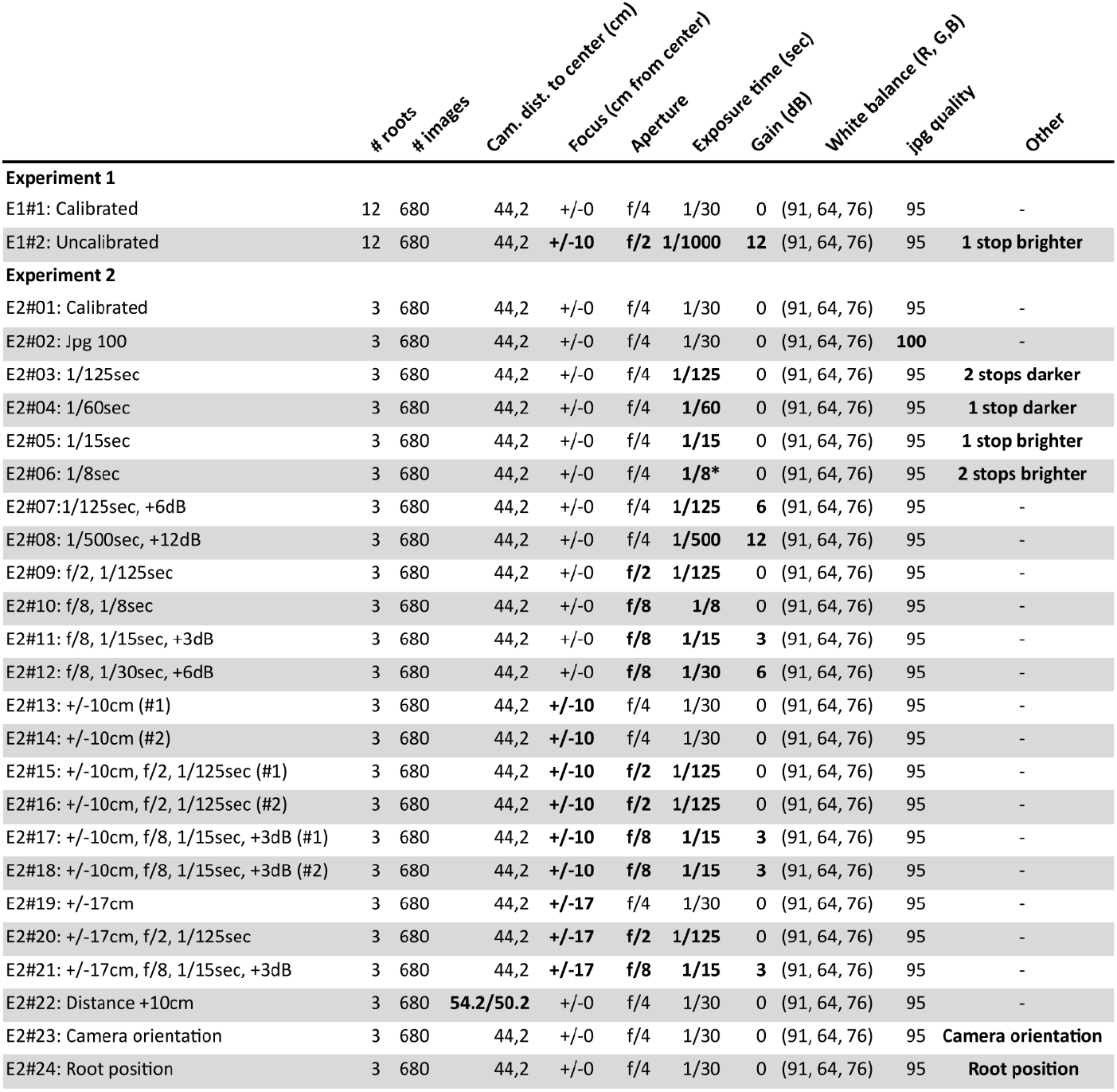
Overview of parameters used in each test of Experiments 1 and 2. Bold characters indicate settings that differ from the calibrated state. The symbol * in test E2#06 indicates that one camera was set to 1/30 sec instead 1/8 sec.

#### 2.4.2 Experiment 2

Experiment 2 consisted of 24 individual tests (E2#01 to E2#24) with three roots (root_3, root_4 and root_9) to further measure the sensitivity of point cloud quality and trait accuracy to individual and combined changes in scanner parameters. The first test (E2#01) acts as our calibrated setup, followed by a series of tests to study the effect of image compression (jpg quality), exposure time and gain. In E2#02 we increased image quality by reducing image compression; in E2#03 to E2#06 we changed exposure time to either under- or overexpose images; and in E2#07 we increased gain to produce grain and at the same time reduced exposure time to keep the same image brightness as in the calibrated setting. In tests E2#13 to E2#21 we changed the camera focus either alone or in combination with narrower or wider DoF. During these tests, we had to reduce exposure time at large aperture (f/2) and increase exposure time and gain at smaller apertures (f/8) to not alter image brightness. Tests E2#22 to E2#24 focused on the geometric setup of scanner and root. In E2#22, we increased the distance between the root and cameras by moving eight cameras back by 10 cm and two back by only 6 cm due to lamp mountings. In E2#23 we turned the cameras’ views by approximately 10° to 15° alternating left and right from the straight view to the scanner center, such that roots were not completely imaged, or lamps were visible in images. Finally, in E2#24 we moved the root 7 cm up and 7 cm to the side. Experiment 2 resulted in a total of 72 scans.

### 2.5 Scanning and Model Reconstruction

To minimize potential effects of other factors on our analysis we performed all scans in the same manner. As such, each root was marked with a black dot on the stem to assure the same placement of each root into the scanner between different tests. Further, the rotating metal frame started in the same position for all scans and revolved over an angle of 335°. Images were taken every 5°, resulting in 680 images per scan. For image acquisition the rotation of the frame was halted. All images were saved in jpg format and have a resolution of 3872 × 2764 pixels. We then computed 3D models for all 24 scans in Experiment 1 and all 72 scans in Experiment 2 with DIRT/3D (COLMAP) as described in Liu, Bonelli (30). In Experiment 2 the reconstruction of root_9 in tests E2#06 and E2#23 failed.

All reconstructed models were then cleaned. Points not belonging to the root were manually cropped and the stem was cropped upward from the marked black dot using CloudCompare v2.12.4 (Kyiv) (40). We further applied a statistical outlier removal (SOR) filter using PyntCloud (41), where for each point the mean distance to its ten nearest neighbors was computed and points with a value larger than the model’s mean plus three times the standard deviation were excluded. This cleaning process resulted in the removal of isolated points and in a visible reduction of noise in the point cloud.

### 2.6 Feature Extraction

We then assessed the effects of calibration and other scanner settings on point cloud quality and accuracy of measured 3D root traits in Experiment 1 and Experiment 2. To assess point cloud quality, we computed the local features curvature, anisotropy, linearity, planarity, and sphericity in each 3D model based on Principal Component Analysis (PCA) using the procedure outlined in Zhang et al. (42). Features were calculated for each point by selecting the *k* nearest neighbors of each point and performing a principal component analysis to calculate the eigenvalues (λ_1_, λ_2_ and λ_3_) of the first, second and third principal components at the local point distribution. Based on the values the shape of the point distribution can be determined. For example, a large λ_1_ compared to λ_2_ and λ_3_ represents a linear point distribution as would be expected in fine roots, while similar values for λ_1_ and λ_2_ compared to a small value for λ_3_ represent a planar distribution as we can expect on the surface of the plant’s stem. As such, we expect the values of linearity and planarity in our root models to decrease with increased noise due to a bad scanner setup.

### 2.7 Statistical Analysis

In Experiment 1 we computed the means of the features curvature, anisotropy, linearity, planarity, and sphericity using *k* equal to 200 in each point cloud before and after application of the SOR filter for models of all tests. We then analyzed the effect of calibration as well as filtering on all five features using a paired t-test. In Experiment 2 we computed the mean values of curvature, anisotropy, linearity, planarity, and sphericity without the SOR filter for all tests (E2#02 – E2#23) and compared them to the calibrated setup (E2#01) as a percentage change.

## 3. Results

### 3.1 Scanner Calibration

#### 3.1.1 Calibration Objectives

The calibration of the root scanner is a critical process that ensures accurate and precise imaging for 3D reconstruction. To achieve this, we established specific calibration goals and defined the necessary actions to meet them (Table 3). Based on these, the first step in the calibration process is setting the position and orientation of the cameras, light sources, and root. This ensures that the root system is well-covered in all images. Once this setup is complete, we focus the cameras on the scanner’s center, where the root will be placed. With the root positioned and the cameras’ focal point aligned in the scanner center, we adjust the DoF by selecting an appropriate aperture setting. Since DoF is a key calibration parameter, the aperture should be set to maximize sharpness across the entire root system, rather than being used to adjust image brightness. However, since light and the camera’s diffraction limit can be limiting factors, the aperture should be kept as large as possible while still maintaining sufficient DoF. After setting the aperture, we adjust the white balance to achieve a natural color representation in the images. Finally, we fine-tune exposure time and gain to ensure adequate brightness without degrading image quality. By following this systematic approach, the scanner can effectively capture detailed imagery of complex root structures, enabling accurate 3D reconstructions and analyses. Each of these steps requires careful manual and visual precision, highlighting the importance of a well-prepared setup for achieving precise calibration, as discussed in the following sections.

**Table 3:**
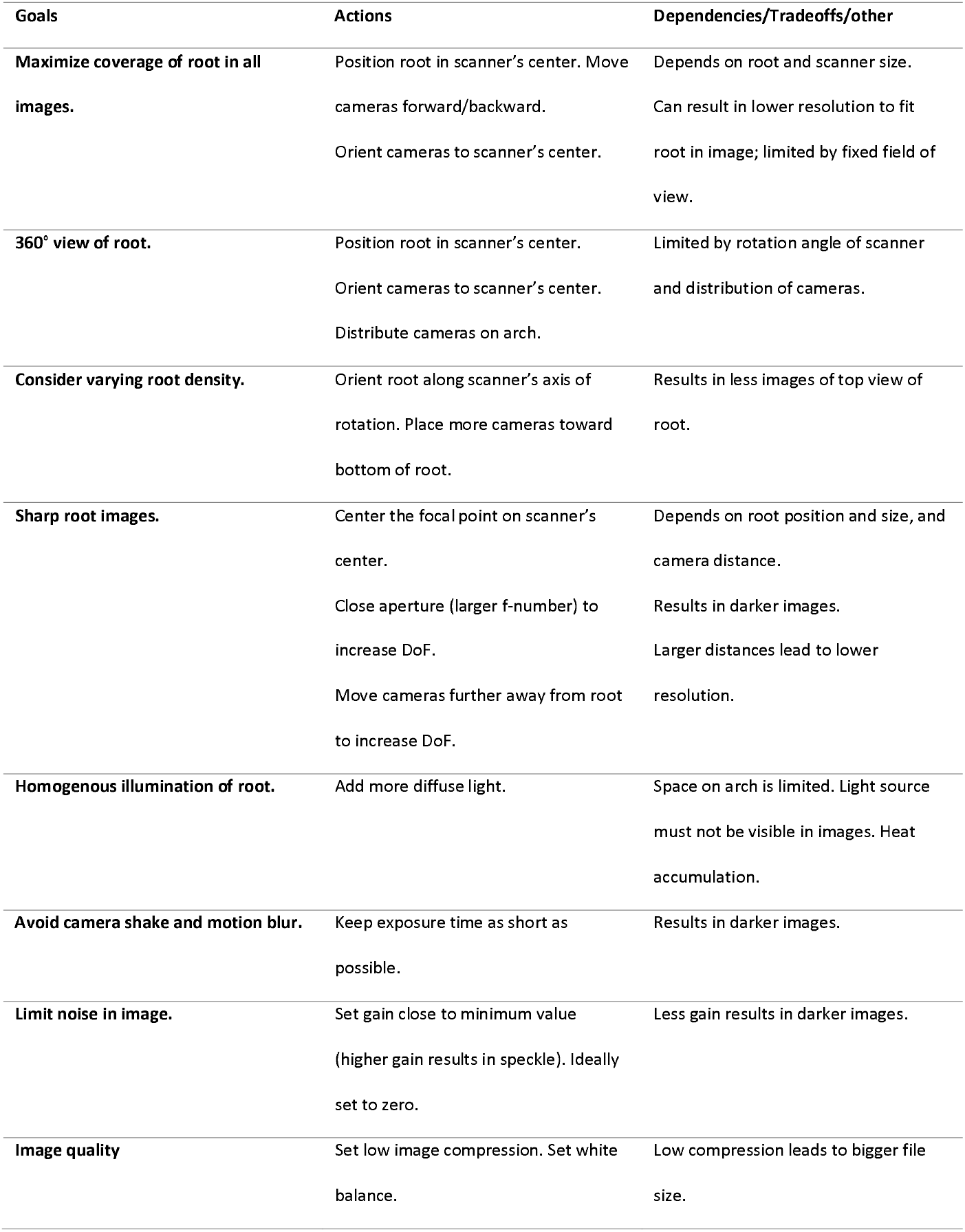
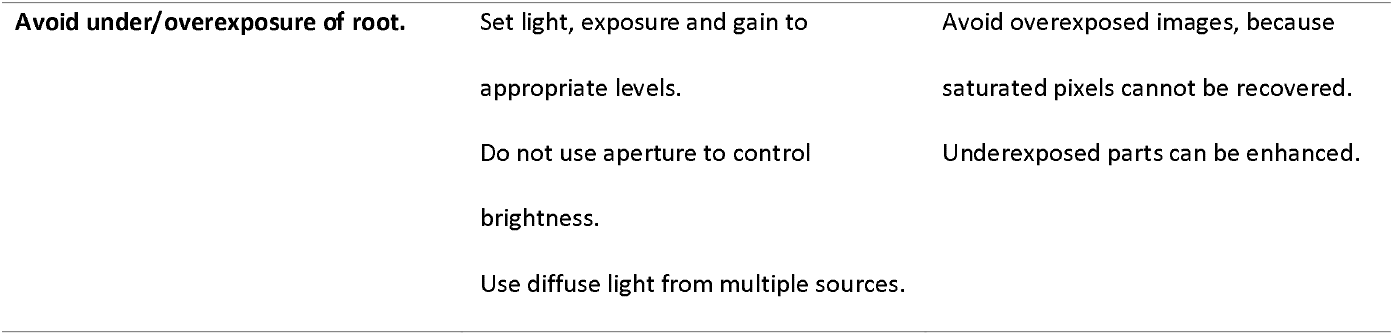
Overview of objectives to achieve high-quality root images.

#### 3.1.2 Scanner Calibration Procedure

##### 3.1.2.1 Pre-calibration

Before initiating the calibration process, baseline adjustments were made to enable initial captures of images and the calibration of camera position and orientation. The power settings of the light sources were set to their maximum value. This preliminary step involved setting rough adjustments of focus, aperture and exposure based on visual inspection of the images to facilitate further adjustment during the calibration process.

##### 3.1.2.2 Root position and orientation

We positioned the root in the center of the scanner such that the distance between the center of the root system and cameras does not change when the camera rig rotates around the root, enabling the use of a constant focusing distance (Figure 2A). We further aligned the stem and the scanner’s axis of rotation, with the stem pointing away from the scanner’s motor and the bottom of the root pointing toward it.

**Figure 2:**
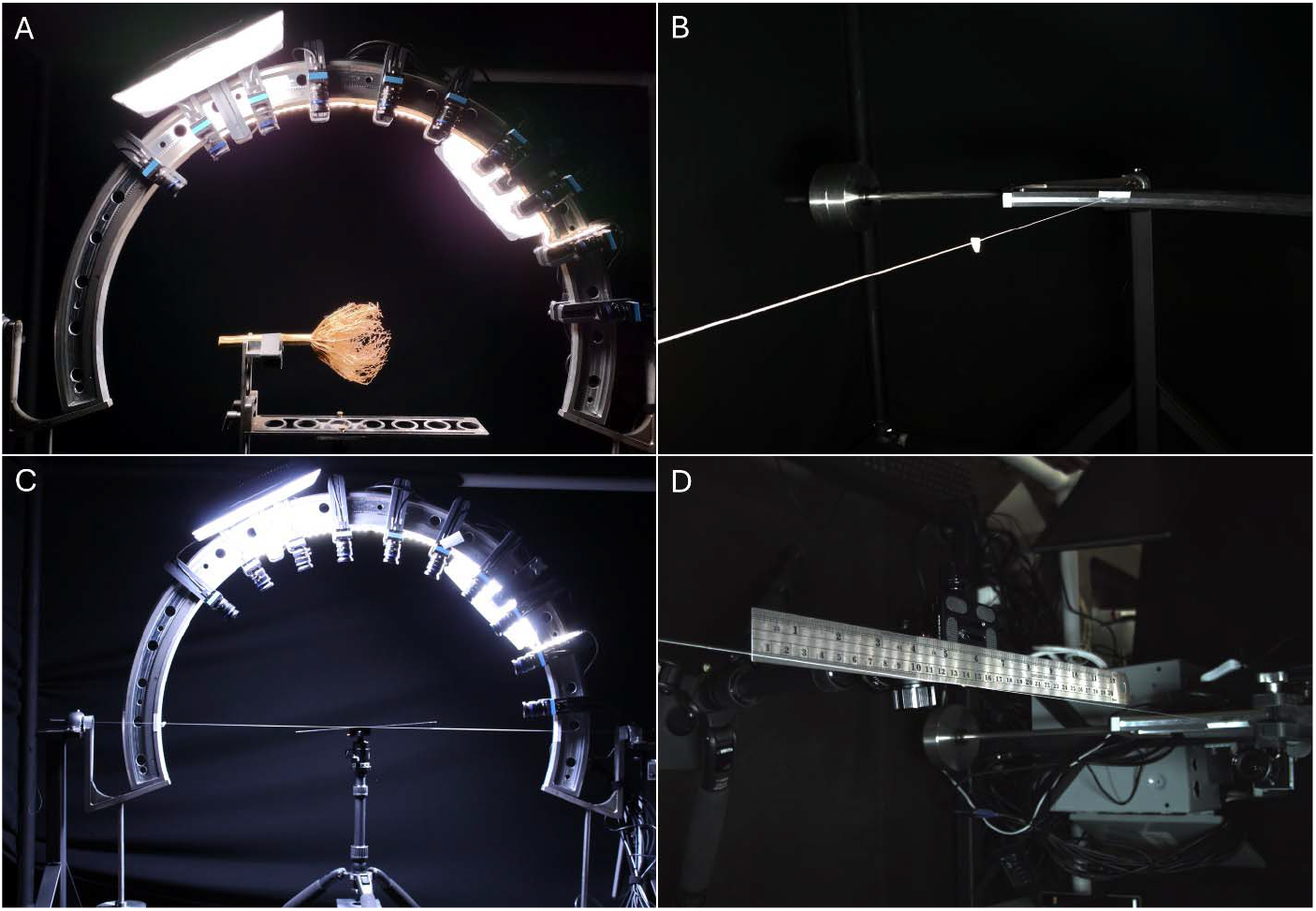
Image of calibrated scanner with maize root and calibration steps. (A)The calibrated scanner with the root in the center. (B) A white marker attached to thread spanned across the center of the scanner as seen from a camera. (C) A ruler was placed in the center of the scanner and tilted to focus each camera. (D) The ruler as seen from a camera.

##### 3.1.2.3 Camera Position

The scanner calibration process started with adjustments to the position of all ten cameras on the arch. The rightmost camera (PiController) was set at an angle of 7.5° from the rotation axis of the camera arch allowing images from the bottom and inside of the root crown (Figure 2A). We uniformly spaced the remaining nine cameras at an angle of 15° between them, resulting in an overall coverage of 135°. All cameras were moved as far forward toward the scanner as was possible on the mounting attached to the arch, resulting in a distance between the scanner center and camera sensors of 44.2cm. Due to thicker roots and fewer fine roots at the top of the root system, we did not place cameras on the left side of the scanner.

##### 3.1.2.4 Camera Orientation

The process of adjusting camera orientation began by positioning a white marker at the center of the scanner, corresponding to the anticipated location of the root (Figure 2B). To achieve this, a white thread was extended across the scanner, spanning from one side to the other along the rotational axis of the scanner arch. Moreover, the midpoint of the thread was identified, and a white marker was affixed to it, ensuring enhanced visibility of the scanner’s center in images captured by all ten cameras. Subsequently, all cameras were aligned to orient the marker as closely as possible to image centers. It should be noted that only adjustments to the camera’s yaw angle (i.e., rotation left or right) were performed due to the fixed mounting of the cameras on the arch. As such, the pitch and roll angles were not adjustable. Throughout this adjustment process, TCam Capture facilitated real-time image monitoring. Consequently, the reorientation of all cameras toward the center of the scanner was achieved.

##### 3.1.2.5 Light Sources

The two light sources were attached to the arch such that they were as far apart without being in the field of view of any of the cameras (Figure 2A). At this point we kept the power of the light source at the maximum setting.

##### 3.1.2.6 Camera Focus

Optimization of camera focus was crucial for maintaining sharp focus of the entire root structure throughout all camera positions. Because the cameras used are not equipped with automatic focus this procedure had to be performed manually and individually for all ten cameras. For focus distance the lens had markings at 0.1m, 0.3m and infinity. As such, this step posed the greatest difficulty and demanded considerable time. To facilitate this process, a ruler was affixed to a camera tripod before being placed inside the scanner (Figure 2C). The ruler was then pointed toward the camera and tilted to produce an angle of 45° between the ruler and the straight line from the scanner’s center to the camera’s center. It was further ensured that the ruler’s center was always aligned with the center of the scanner during the entire process. The aperture of the camera was then set to the lowest f-number (f/1.6) by turning the aperture ring of the lens to create a shallow DoF. An image was captured using TCam Capture, and the focus point was determined based on the sharpness of the ruler’s lines and numbers in the image (Figure 2D). The focus was then adjusted to focus on the center of the ruler by turning the focus ring accordingly. If the focus was found to be too close to the camera, the focus distance was increased; conversely, if the focus was too far from the camera, it was decreased. Another image was captured using the TCam Capture tool to verify the accuracy of the focus. This process was repeated until optimal results were achieved for all cameras.

##### 3.1.2.7 Aperture

Ensuring an optimal DoF was paramount to maintain sharpness across the entire root system. We utilized the same setup with a ruler from the previous calibration step to facilitate DoF assessment (Figure 2D). Visual inspection of the sharpness of the ruler’s lines and numbers in images served as our guide in selecting the ideal aperture setting. Our criterion for achieving sufficient DoF encompassed the sharpness of all markings on the 30 cm long ruler. The aperture range of our camera lenses spanned from f/1.6 (large iris) to f/16 (small iris). We initiated the process with an aperture setting of f/2 as our starting point, progressively increasing the f-number using only settings marked on the camera lenses (i.e., f/2, f/2.8, f/4, f/8) until the desired DoF was attained. These settings facilitated straightforward calculations of exposure times needed to compensate for the smaller iris by doubling exposure time between f/2 and f/2.8, f/2.8 and f/4, and quadrupling exposure time between f/4 and f/8, ensuring consistent image brightness across different aperture settings.

Compensating for darker images resulting from smaller apertures necessitated a corresponding increase in exposure time. Given the inherent trade-off between DoF and image brightness associated with aperture adjustment, we selected f/4 as the largest aperture that would still yield sharp representations of all parts of the root system.

##### 3.1.2.8 White balance

To also ensure naturally looking images the white balance of the camera was calibrated. To obtain the value for white balance we placed a white sheet of paper in front of a camera and used TCam Capture’s feature to automatically adjust white balance. This was repeated for each camera and the detected values for red, green and blue were averaged across all cameras, resulting in the values 91, 64 and 76 for the red, green and blue channels, respectively. These values were used in TCam Capture to set the white balance.

##### 3.1.2.9 Exposure time

As previously mentioned, it is generally imperative to maintain the exposure time as short as possible to mitigate camera shake and motion blur. Due to the non-moving root and the static position of the cameras during exposure we expected that these would not be a major concern. However, we observed that thin and long roots vibrated. In either case of short and long exposure time this would lead to inaccuracies during 3D reconstruction and neither short nor long exposure times would be able to alleviate its effect. In this setup, an exposure of 1/30 sec resulted in images with sufficient brightness, while not overexposing any parts of the root system.

##### 3.1.2.10 Gain

Finally, the gain was adjusted. After aperture, exposure time and illumination, an increase in gain is the last possibility to enhance image brightness. Nevertheless, high gain also leads to increased noise levels in the form of speckle within the image. Therefore, efforts were made to maintain the gain at the lowest level. Since the exposure time of 1/30 sec seemed to suffice for image brightness, the gain was set to its lowest value of 0 dB (TCam Capture setting of 100).

### 3.2 Experiment 1

We used all 12 roots to compare the calibrated scanner setup versus an uncalibrated scanner setup, resulting in 24 root models. All models were manually cleaned and cropped, and outliers were removed with the SOR filter. For all models the five features and number of points were calculated before and after applying the SOR filter.

#### 3.2.1 Quality assessment

The effects of calibrating the scanner and cameras on the images of the roots can be easily observed by visual inspection of the images. In the image of the calibrated scanner the complete root system is well exposed, i.e. all roots are sufficiently bright and none are overexposed (Figure 3, top). Further all parts in the front and back of the root remain sharp, and details such as fine root can be seen as well as the resolution of the image allows. In contrast, the root in the uncalibrated image is overexposed due to the combined effects of a large aperture and high gain; the roots that are closer to the camera are blurred due to the incorrect focus and large aperture, and speckle is visible due to the high gain.

**Figure 3:**
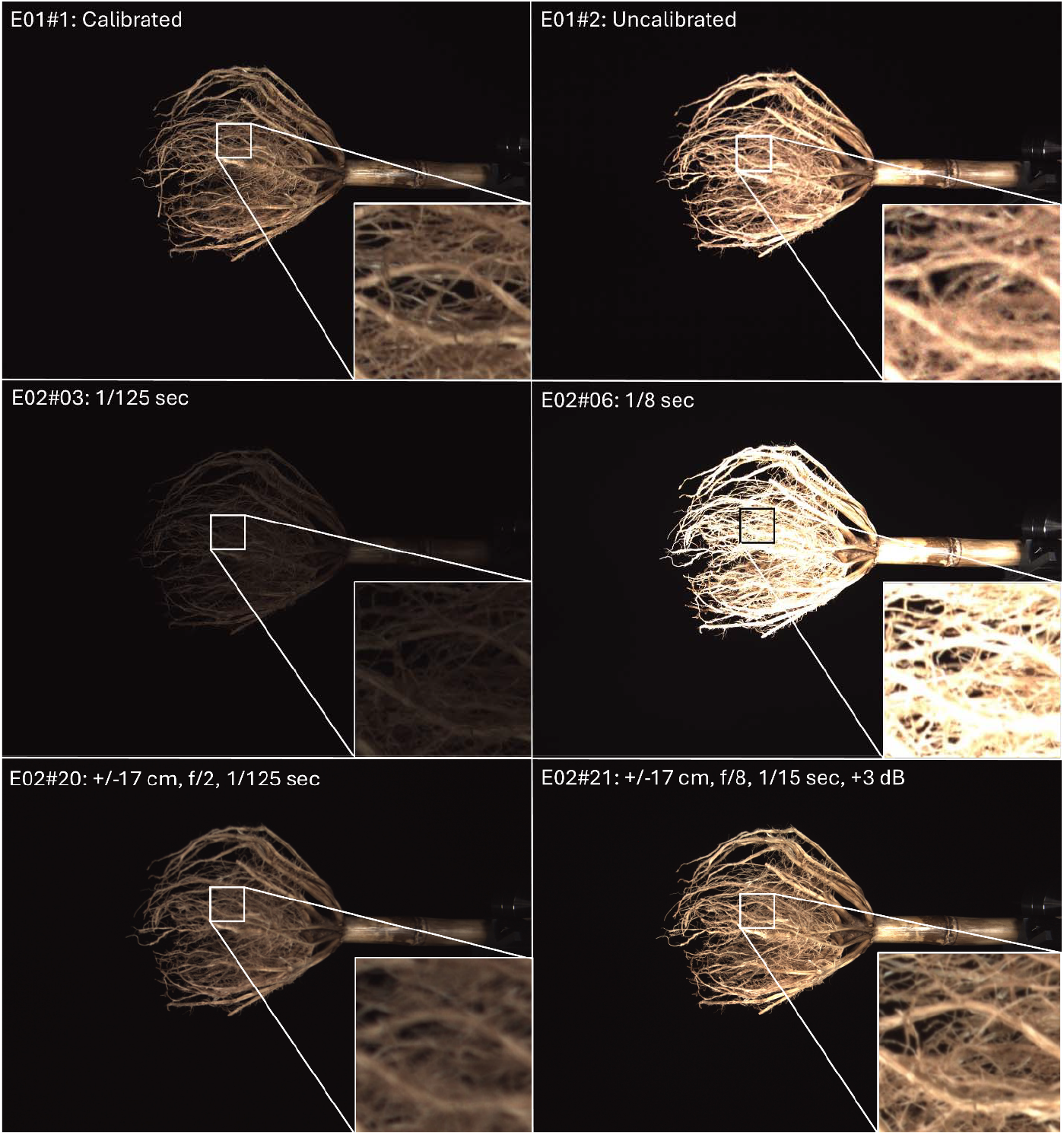
Images of root_9 from six different tests. The boxes zoom into a small part of the root crown in each image to highlight differences coming from different scanner setups. The top row shows the calibrated and uncalibrated setup, the middle row the effects of image brightness (due to exposure time), and the bottom row the combined effects of focus and aperture.

A visual presentation of a 3D model generated with COLMAP after scanner calibration and its linear, planar, and spherical features are given in Figure 4. A comparison of the 3D point cloud with the image corresponding to the same viewing angle shows accurate resemblance between the overall root structures as well as fine roots. Generally, it can be observed though that fine roots appear thicker in the 3D model than in the image. The visualization of linearity, planarity, and sphericity shows differences between separate parts of the root. Due to the large surface area the stem has the largest values in planarity, while linearity and sphericity are low. Some smaller regions on the stem show smaller values of planarity and higher values of sphericity. Larger values of planarity can be further seen at the thickest roots. Linear features can only be detected in fine roots within the root crown if they are sufficiently thin. Overall, only a few points exhibit this high linearity. Regarding sphericity no visible points exhibit a large value, but many points corresponding to thicker roots in the crown root show a moderate sphericity around a value of 0.5.

**Figure 4:**
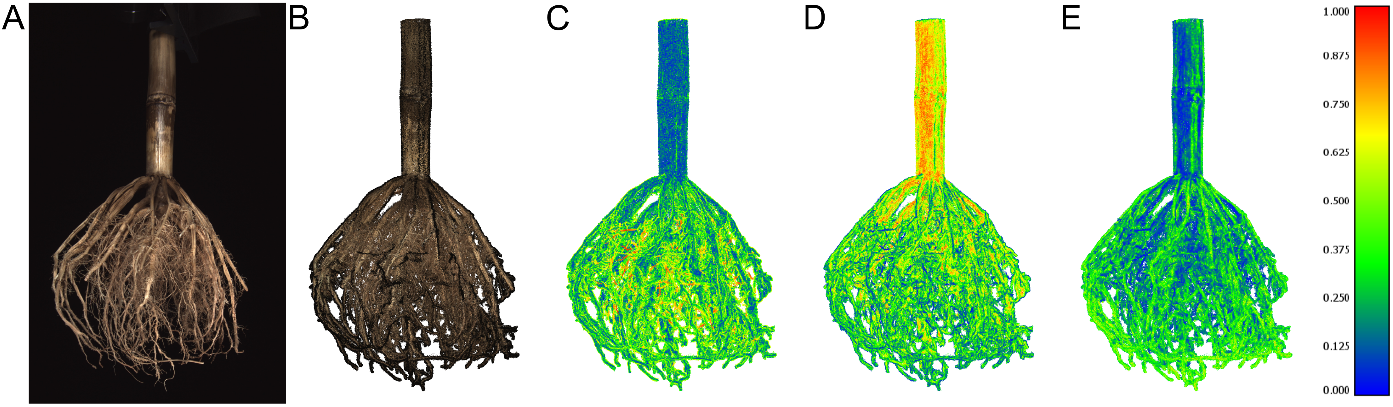
Image and 3D models with original root color and computed features for root sample root_9. (A) Cropped jpg image of root taken during the scanning process with calibrated scanner. (B) 3D point cloud of root after reconstruction with COLMAP an after cleaning and filtering, where points correspond to 3D measurements of the root. (C-E) Features illustrating the local distribution of points using 200 nearest neighbors (k=200), where (C) depicts linearity, (D) planarity and (E) sphericity on a scale from 0 to 1. High values of linearity, planarity or sphericity are thus shown in red, whereas low values are shown in blue.

The effects of calibration and filtering on the reconstructed models are visualized by the noise present on the stem cross-section in Figure 5. In both the uncalibrated and calibrated setups several isolated outliers are present because of mismatched image pixels during the 3D reconstruction. This type of noise was successfully removed in both cases by application of the SOR filter. Nonetheless, the root model from the uncalibrated scanner exhibits a rougher stem surface with more noise than the root model from the calibrated scan. This characteristic even prevails after the application of the SOR filter. As such, we see that the root model from the calibrated scan together with filtering results in the most accurate representation of the planar stem surface.

**Figure 5:**
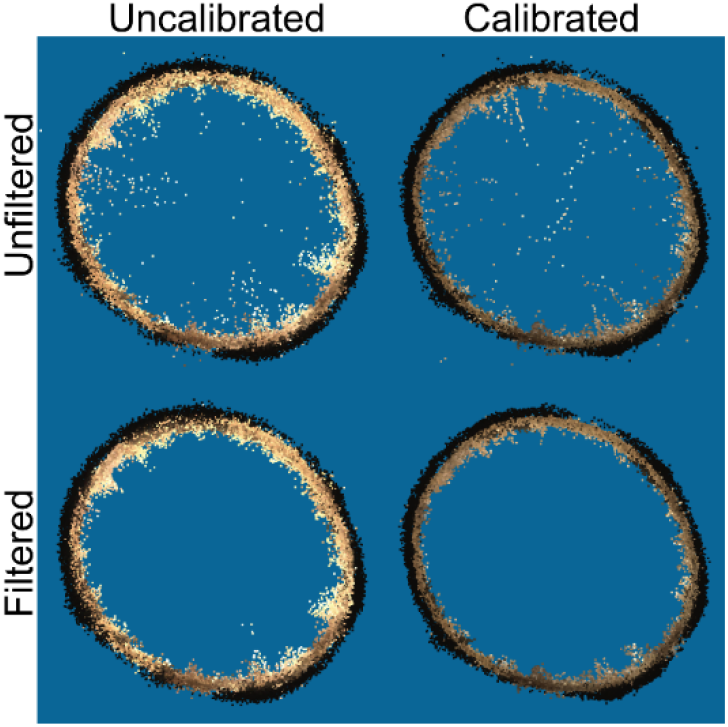
Effect of calibration and filtering on point cloud quality shown at the cross-section of the stem. Cross-sections of stems were cropped above the root crown from point clouds of sample root_9 and have a width of approximately 1 cm. The top-left cross-section was taken from the uncalibrated and unfiltered, top-right from calibrated and unfiltered, bottom left from uncalibrated and filtered, and the bottom right from the calibrated and filtered model.

For the calculation of the five features, we used a value of 200 for the number of neighbors, *k*. Our choice was based on an assessment of the mean curvature, mean anisotropy, mean linearity, mean planarity, and mean sphericity using different values for *k* (10, 25, 50, 100, 200, 300, 400) in all twelve models from test E1#1. The logic for choosing a value of 200 was therefore that planarity converges to a relatively stable value at *k* equal to 200 (see Figure 6). All features were computed with PyntCloud (41).

**Figure 6:**
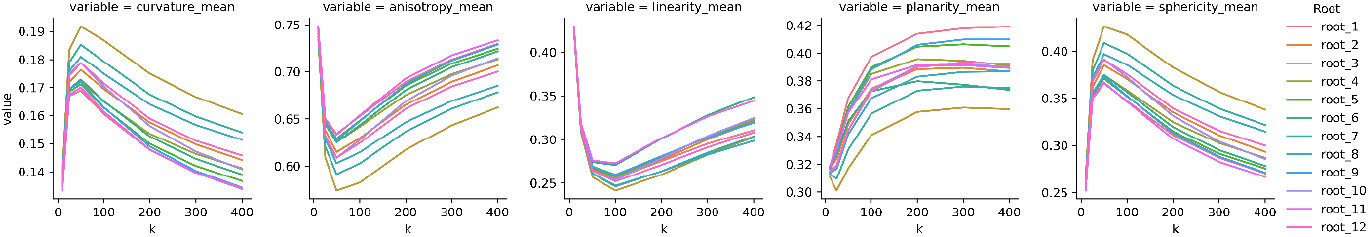
Mean of the five features (curvature, anisotropy, linearity, planarity, and sphericity) for 12 root models as a function of k (number of nearest neighbors). Each line depicts the mean value of a single root model. Features were computed on models from the calibrated scanner setup in test E1#1 and before the application of the SOR filter.

Computation of the five features enabled us to quantify differences in local point distributions between 3D models. Figure 7 shows that local point distributions of the whole 3D model change because of the scanner calibration. Overall, we observed an increase in linearity, planarity and consequently anisotropy, and a decrease in curvature and sphericity. We further observed the same effect of scanner calibration on local point distribution in all twelve roots (see Figure 8). In models from the calibrated scan without SOR filter the mean curvature across all 12 roots decreased by 7.1% from 0.169 to 0.157 (p<0.001), mean anisotropy increased by 5.2% from 0.634 to 0.667 (p<0.001), mean linearity increased by 6.3% from 0.261 to 0.278 (p<0.001), planarity increased by 4.5% from 0.373 to 0.389 p<0.001) and sphericity decreased by 9.1% from 0.366 to 0.333 (p<0.001) compared to models from uncalibrated scans (see Table 4). Similarly, we saw a mean decrease in the number of points by 3.2% from 5,702,920 to 5,519,870.4 (p<0.01).

**Table 4:**
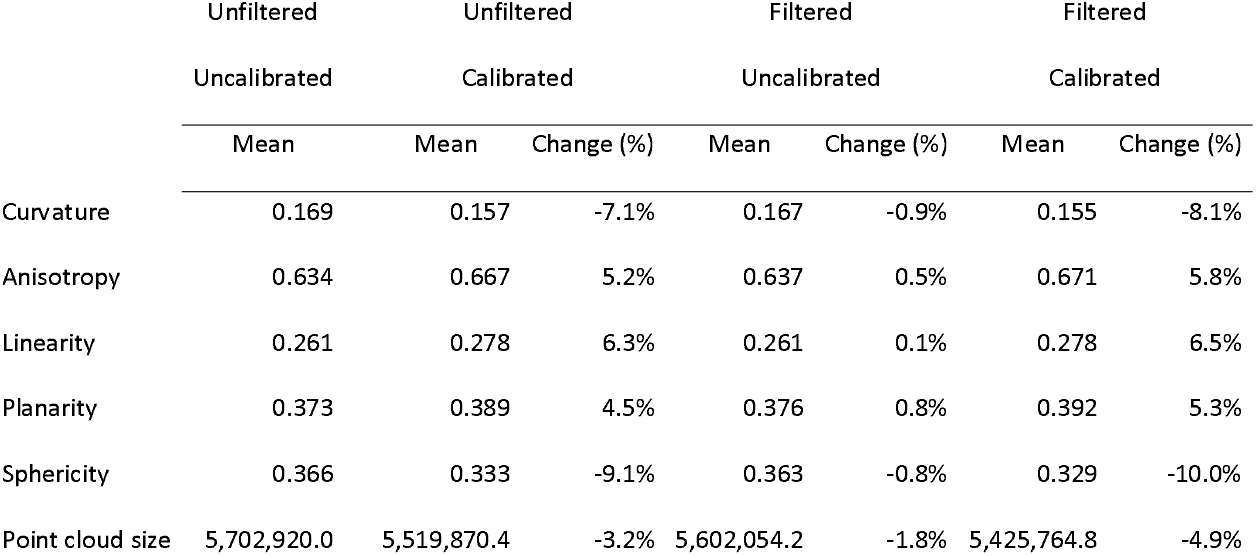
Mean values for curvature, anisotropy, linearity, planarity, sphericity, and point cloud size over all twelve roots. Mean values are shown for uncalibrated-unfiltered, calibrated-unfiltered, uncalibrated-filtered, and calibrated-filtered point clouds. Change is computed with respect to uncalibrated-unfiltered root models.

**Figure 7:**
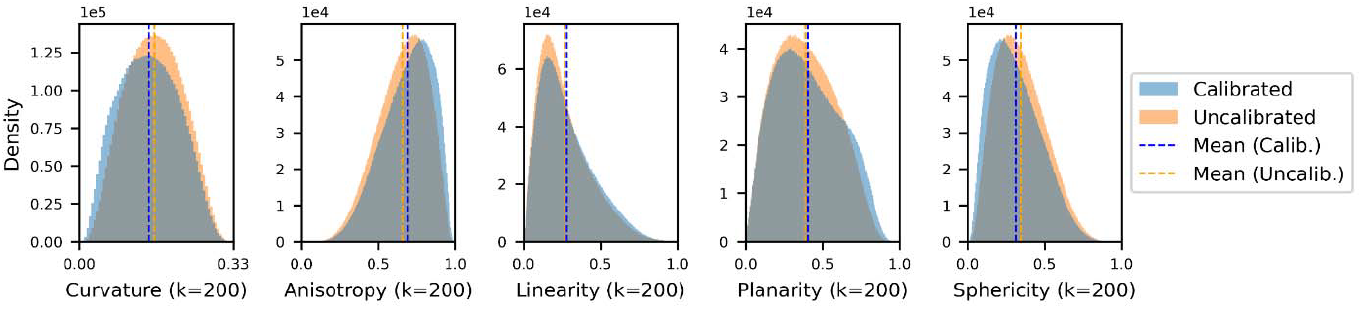
Histograms showing the distribution and mean values of curvature, anisotropy, linearity, planarity, and sphericity of sample root_9 before and after scanner calibration. Features were computed before the application of the SOR filter and with k=200. The width of each bin is 0.005.

**Figure 8:**
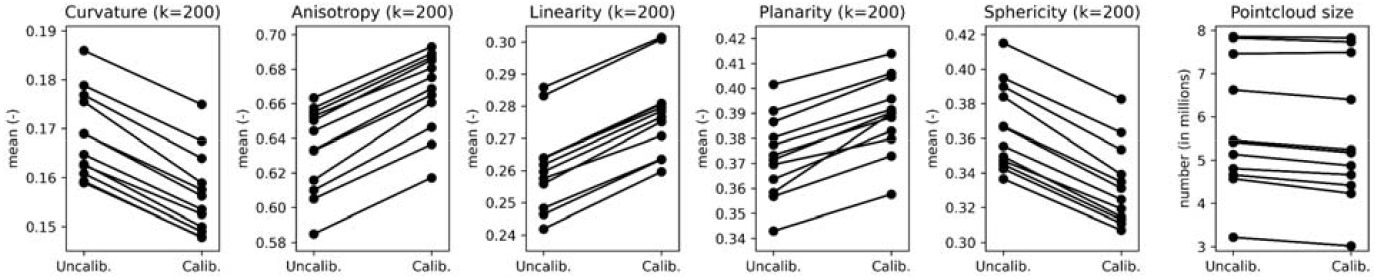
Effect of calibration on curvature, anisotropy, linearity, planarity, sphericity, and point cloud size for paired samples. Values on the left and right in each plot correspond to measurements of point clouds from uncalibrated and calibrated scans, respectively. For curvature, anisotropy, linearity, planarity, and sphericity each filled circle represents its mean calculated for a single point cloud. Point cloud size is shown as the total number of points in each point cloud. All values were computed before the application of the SOR filter. The lines connecting circles indicate corresponding pairs of models before and after calibration.

We further tested which effect filtering has on the local point distribution by comparing features before and after the application of the SOR filter in the uncalibrated scan (see Figure 9). In models after SOR filtering the mean curvature decreased by 0.9% from 0.169 to 0.167 (p<0.001), mean anisotropy increased by 0.5% from 0.634 to 0.637 (p<0.001), mean linearity increased only by 0.1% (p<0.01), planarity increased by 0.8% from 0.373 to 0.376 (p<0.001) and sphericity decreased by 0.8% from 0.366 to 0.363 (p<0.001) compared to models without filtering (see Table 4). Similarly, we saw a decrease in the mean number of points by 1.8% from 5,702,920 to 5,425,764.8 (p<0.001). Further, we observed that the sum of the individual effects of calibration and filtering at large adds up to the same amount as the effect of calibration combined with filtering (Table 4).

**Figure 9:**
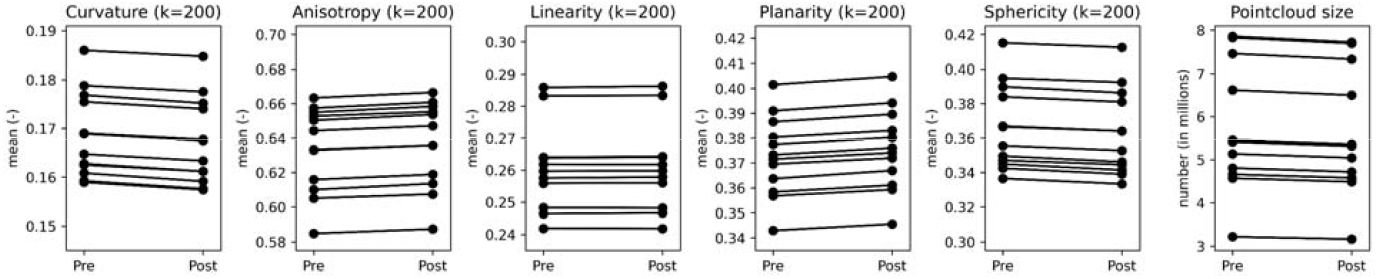
Effect of SOR filter on curvature, anisotropy, linearity, planarity, sphericity, and point cloud size for paired samples. Values on the left and right in each plot correspond to measurements of point clouds before and after the SOR filter, respectively. For curvature, anisotropy, linearity, planarity, and sphericity each filled circle represents its mean calculated for a single point cloud. Point cloud size is shown as the total number of points in each point cloud. All values were computed for the uncalibrated scanner setup. The lines connecting circles indicate corresponding pairs of models before and after the SOR filter.

### 3.3 Experiment 2

In Experiment 2 we performed a total of 24 tests using three models (only two models in tests E2#06 and E2#23). All resulting models were manually cleaned and cropped. Next the five features, number of points, and root traits were computed without the application of the SOR filter.

#### 3.3.1 Quality assessment

The individual adjustments to the scanner setup can have a significant impact on the quality of the images taken. In the highly underexposed images of test E02#03 with an exposure time of 1/125 sec the image is just bright enough that the root can be seen, while in the overexposed images of test E02#06 with an exposure time of 1/8 sec most parts of the root are severely overexposed, such that they appear as plain white and hence fewer features of the fine roots are visible (Figure 3, middle). In images from test E02#20, the use of a large aperture (f/2) resulted in bad focus and shallow DoF, causing most details in fine roots to appear too blurred to discern. In contrast, the issue is largely resolved in images from test E02#21, where a small aperture (f/8) resulted in a greater DoF (Figure 3, bottom)

The effect of changes to individual scanner settings or groups of camera settings on the five features and point cloud size is shown in Figure 10. Overall, most tests had little to no effect (under 1-2%) on curvature, anisotropy, linearity, planarity, sphericity or point cloud size. The largest decrease and the largest increase in curvature relative to the calibrated scan were in test E2#24 with modified root position (−5.3%) and test E2#20 with an unfocused camera and large aperture (24.4%), respectively. The largest decreases in anisotropy (−16.2%), linearity (−13.6%) and planarity (−18.0%) were also in test E2#20 with an unfocused and large aperture setting. And the largest increases in anisotropy (3.5%), linearity (4.1%) and planarity (3.0%) were in test E2#24 with modified root position. The largest decrease in point cloud size was in test E2#20 with an increased camera-to-root distance (3.3%) and the largest increase was in test E2#16 with an unfocused camera and large aperture. Despite the large effect in some tests, only 8 out of 23 tests had deviations in either feature values or point cloud size from the calibrated setup that are larger than five percent.

**Figure 10:**
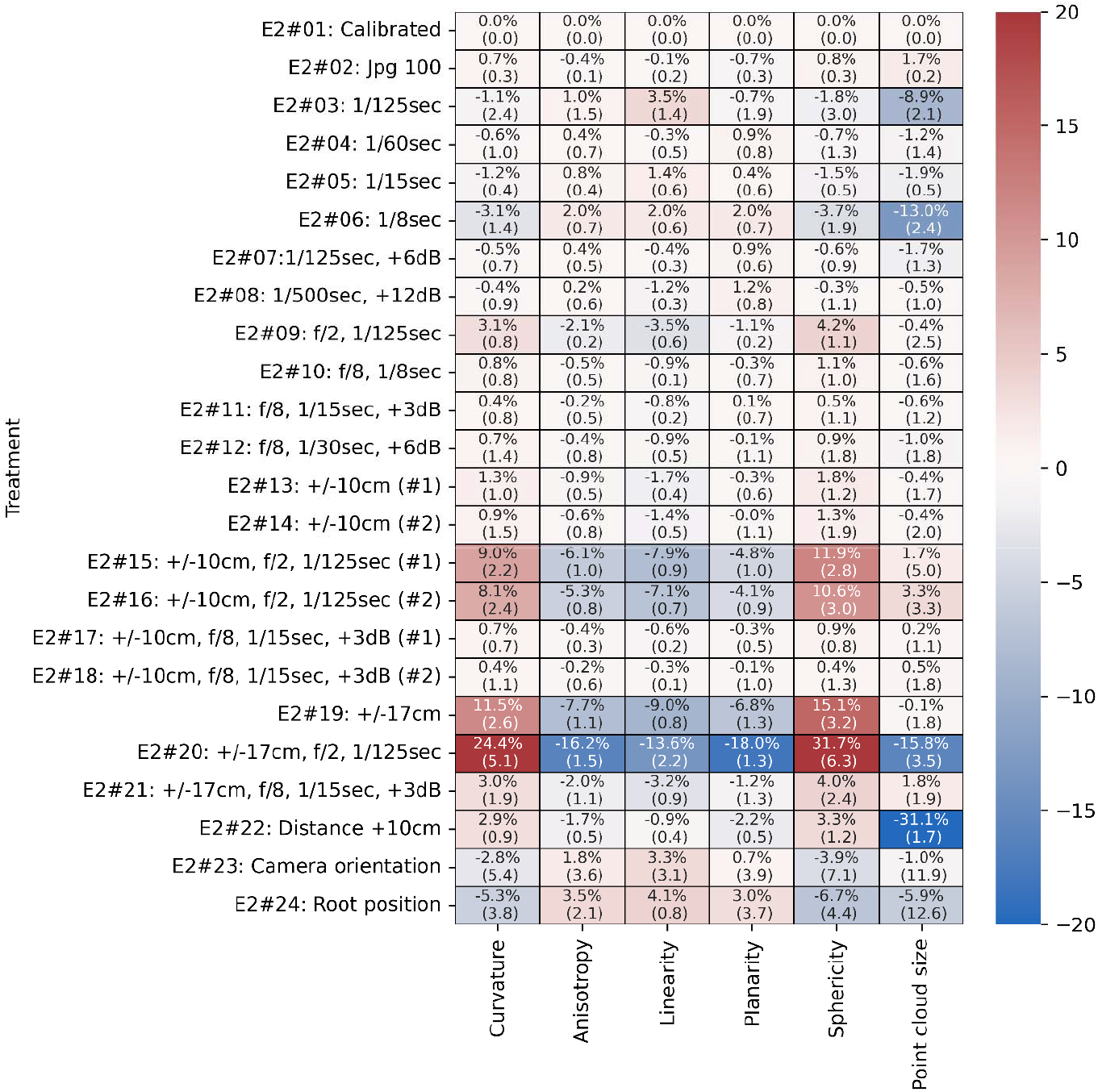
Heatmap of results from Experiment 2 showing effects of individual scanner settings on curvature, anisotropy, linearity, planarity, sphericity, and point cloud size. The first number represents the mean change in percent with respect to the calibrated scanner setup (E2#01) and the number in parentheses represents its standard deviation. Red colors indicate an increase and blue a decrease.

We observed noteworthy effects in tests related to very short or long exposure times, large apertures, unfocused setups and altered scanner-root geometries. Setups with short (E2#03) and long (E2#06) exposure times, i.e. under- and overexposed images, resulted in a smaller point cloud size but had only a small effect on geometric features. Increasing the aperture to f/2 (E2#09) resulted in slightly worse geometric features but reducing it to f/8 (E2#10) only resulted in small differences under 1.1% compared to the calibrated scans, indicating that the narrower DoF caused by the larger aperture of f/2 resulted in a decrease in model quality. An aperture of f/2 and the focal point being 10cm off-center (E2#15 and E2#16) resulted in worsening of all five features but did not affect point cloud size. Contrary to that, the focal point being 10cm off-center with our standard aperture of f/4 (E2#13 and E2#14) and a small aperture of f/8 (E2#17 and E2#18) showed no effect on any feature or the point cloud size. Further, moving the focal point 17cm off-center with a large aperture of f/2 (E2#20) resulted in the largest increases in curvature and sphericity, the largest decreases in anisotropy, linearity, and planarity, and the second largest decrease in point cloud size over all tests. This effect was reduced by using a smaller aperture of f/4 (E2#19), and almost completely removed by using an aperture of f/8 (E2#21). These results show that increased distance of the focal point from the scanner center, the root thus being unfocused, and shallow DoF due to larger apertures reduce model quality. Their combined effect of unfocused setup and shallow DoF amplifies the decrease even further. The effect of an unfocused camera can, to a large extent, be compensated for by the large DoF stemming from a small aperture and vice versa, the effect of a shallow DoF can be compensated for by accurate focusing on the scanner center. Increasing the distance between cameras and the root resulted in the largest observed reduction in point cloud size and moving the root off-center on average improved features and reduced point cloud size, but the effect was driven by one root only.

## 4 Discussion

### 4.1 Summary of findings

In this study, we created a methodical calibration pipeline to improve a multi-camera rig used for 3D reconstruction of plant roots. We then tested how camera calibration affected the quality of 3D reconstructions of maize root crowns, paying special attention to camera settings like focus, aperture, exposure time, and gain. Our results demonstrate that proper calibration of these settings significantly enhances the quality of reconstructed 3D models. Specifically, misalignment of the focal plane and shallow DoF led to increased noise and loss of fine root structures. Our results show how important it is to properly calibrate a scanner in order to improve the quality of a point cloud, lower noise, and make root trait extraction more reliable.

The results from Experiment 1 that using a properly calibrated scanner greatly improves the accuracy of 3D reconstructions. This was shown by higher linearity, planarity, and anisotropy values, as well as lower curvature and sphericity values. These improvements indicate a more structured and less noisy point cloud, which is critical for later steps of the analysis like root skeletonization and trait extraction. Furthermore, filtering using the SOR method, which is useful to remove outliers, had a relatively small effect compared to the benefits of proper scanner calibration. This suggests that reducing noise at the image acquisition stage is more effective than relying solely on post-processing steps.

Experiment 2 provided more detailed insights into the sensitivity of 3D reconstruction quality to individual camera settings. We observed that small deviations from the calibrated setup, such as slight changes in exposure time or gain, had minimal impact on point cloud quality. However, more substantial deviations, in particularly poor focus combined with a shallow DoF (achieved by using a large aperture), resulted in a severe decline in model quality. Interestingly, our results also demonstrated that increasing the DoF (by reducing the aperture size) could partially compensate for slight misfocusing. However, this trade-off required adjusting exposure time and gain to maintain proper image brightness. These findings underscore the importance of carefully selecting camera settings to optimize both sharpness and brightness while minimizing noise.

### 4.2 Comparison to other fields

The role of camera settings in 3D reconstruction is well understood in photogrammetry, yet its systematic evaluation in plant phenotyping remains limited. While proper camera calibration has been studied in forestry (43), archaeology (44, 45), and in the geosciences (46, 47), plant phenotyping presents unique challenges. Unlike forestry and remote sensing, where large-scale features dominate and motion blur is a primary concern due to the fast movement of camera, root phenotyping deals with small, highly occluded structures, making DoF and focus calibration critical. Studies in archaeology (48) and small object reconstruction (49, 50) also emphasize DoF as a major limitation in close-range photogrammetry. However, these studies often suggest focus stacking or tilt-shift lenses as solutions, which may not be feasible for high-throughput plant phenotyping due to time constraints and system complexity.

Our results align with recommendations from these fields, particularly in emphasizing manual focus control, optimizing the exposure triangle (gain/ISO, exposure time, aperture), and maintaining consistency in scanning geometry. However, root phenotyping requires additional considerations, such as accounting for organic structures with varying surface properties and fine root details, which are more sensitive to poor calibration than rigid objects in archaeological photogrammetry.

### 4.3 Limitations and Future Work

Despite these findings, some limitations in our research remain. First, our study did not evaluate the effects of intrinsic and extrinsic calibration (e.g., focal length, lens distortion, and camera poses), as this was handled automatically by COLMAP. Future work could explore how manual intrinsic and extrinsic calibration using calibration boards compares with automated methods in terms of reconstruction accuracy.

A second limitation of this study is that we did not explicitly test the effects of diffraction at very small apertures (e.g., f/11–f/16), which could reduce sharpness due to light wave interference (50, 51). While our findings suggest f/4 to f/8 as an optimal range, future work could systematically analyze the diffraction limit in plant phenotyping setups. Additionally, light distribution was not fully explored, which could introduce artifacts in 3D reconstructions, a factor that may also influence phenotyping workflows. Future research could test controlled lighting environments to further optimize 3D scanning conditions.

Additionally, while our results demonstrate that a calibrated setup produces more reliable point clouds, further validation with different plant species and environmental conditions is needed to assess the generalizability of our findings. Moreover, variations in point cloud density might influence k-nearest neighbor calculations used for feature extraction, which should be considered in future studies. Finally, this study demonstrates that scanner calibration significantly improves 3D model quality, but we did not explicitly assess its impact on root trait measurements, such as root length, diameter, or volume, which should be investigated in future studies to fully understand the biological implications of improperly setup calibrated camera setups.

## 5 Conclusion

In this study, we developed and validated a camera calibration pipeline aimed at improving the accuracy and reproducibility of 3D root phenotyping. Our results show that careful adjustment of focus, aperture, and exposure settings significantly improves model accuracy, and that calibration has a stronger effect on point cloud quality than post-processing filters alone. Furthermore, we identified the most critical factors affecting reconstruction accuracy, with poor focus and a shallow DoF being the most detrimental. Importantly, we demonstrated that increasing DoF can mitigate the effects of slight misfocusing, providing practical guidelines for optimizing camera settings in future studies.

Our findings contribute to the broader effort of improving transparency and reproducibility in plant phenotyping. By providing a structured calibration workflow, we aim to assist researchers in setting up robust imaging pipelines that produce high-quality 3D models. Future research should explore additional calibration factors, such as intrinsic and extrinsic parameter refinement and different lighting conditions, to further enhance the robustness of 3D root phenotyping methodologies. Ultimately, improved imaging practices will facilitate more precise quantification of root traits, advancing our understanding of plant growth and resilience in diverse environments.

## 6 Declarations

### 6.1 Ethics approval and consent to participate

Not applicable.

### 6.2 Consent for publication

Not applicable.

### 6.3 Availability of data and materials

All data used in the manuscript is available in the BioImage Archive under the DOI: 10.6019/S-BIAD1690

### 6.4 Competing interests

The authors declare that they have no competing interests.

### 6.5 Funding

The research was supported by the NSF CAREER Award No. 2329282 and by the USDOE ARPA-E ROOTS Award Number DE-AR0000821 to A.B. Any opinions, findings, and conclusions or recommendations expressed in this material are those of the authors and do not necessarily reflect the views of the National Science Foundation or the US Department of Energy.

### 6.6 Authors’ contributions

P.P., S.L. and A.B. conceived the calibration pipeline, experiments, algorithms, and data analysis. P.P. and A.B. wrote the original manuscript. P.P. collected data, implemented algorithms and performed data analysis. All authors reviewed and edited the manuscript. AB conceived the project and acquired funding.

## 6.7 Acknowledgements

The work was conducted while transitioning institutions for AB and SL (University of Georgia to University of Arizona).

## Notes

https://doi.org/10.6019/S-BIAD1690

### Competing Interest Statement

The authors have declared no competing interest.

