## Supplementary material for "Optimizing root phenotyping: Assessing the impact of camera calibration on 3D root reconstruction": feature_extract.py

```
# Based on:
#
# Z. Zhang, W. Sun, X. Min, T. Wang, W. Lu and G. Zhai, 
# "No-Reference Quality Assessment for 3D Colored Point Cloud and Mesh Models," 
# in IEEE Transactions on Circuits and Systems for Video Technology, 
# vol. 32, no. 11, pp. 7618-7631, Nov. 2022, doi: 10.1109/TCSVT.2022.3186894.
# https://github.com/zzc-1998/NR-3DQA

import numpy as np
import pandas as pd
from skimage import color
from nss_functions import * # https://github.com/zzc-1998/NR-3DQA
from pyntcloud import PyntCloud
import os 
from time import sleep
import argparse

def get_feature_vector(objpath, k):  
    #load colored point cloud
    print("Begin loading point cloud")
    cloud = PyntCloud.from_file(objpath)

    #begin geometry projection
    print("Begin geometry feature extraction.")
    k_neighbors = cloud.get_neighbors(k=k)
    ev = cloud.add_scalar_field("eigen_values", k_neighbors=k_neighbors)
    cloud.add_scalar_field("curvature", ev=ev)
    cloud.add_scalar_field("anisotropy",ev=ev)
    cloud.add_scalar_field("linearity",ev=ev)
    cloud.add_scalar_field("planarity",ev=ev)
    cloud.add_scalar_field("sphericity",ev=ev)
    kplusone = k+1
    curvature = cloud.points['curvature({})'.format(kplusone)].to_numpy()
    anisotropy = cloud.points['anisotropy({})'.format(kplusone)].to_numpy()
    linearity = cloud.points['linearity({})'.format(kplusone)].to_numpy()
    planarity = cloud.points['planarity({})'.format(kplusone)].to_numpy()
    sphericity = cloud.points['sphericity({})'.format(kplusone)].to_numpy()


    #begin color projection
    print("Begin color feature extraction.")
    rgb_color = cloud.points[['red','green','blue']].to_numpy()/255
    lab_color = color.rgb2lab(rgb_color)
    l = lab_color[:,0]
    a = lab_color[:,1]
    b = lab_color[:,2]


    print("Begin NSS parameters estimation.")
    # compute nss parameters
    nss_params = []
    # compute color nss features
    for tmp in [l,a,b]:
        params = get_color_nss_param(tmp)
        #flatten the feature vector
        nss_params = nss_params + [i for item in params for i in item]
    # compute geomerty nss features
    for tmp in [curvature,anisotropy,linearity,planarity,sphericity]:
        params = get_geometry_nss_param(tmp)
        #flatten the feature vector
        nss_params = nss_params + [i for item in params for i in item]
    return nss_params


if __name__ == '__main__':
    #demo
    #input_path = "D:/E1_1_Calibrated/root_1_good/root_1_good_clean.ply"
    #output_path = "result_feature_extraction.csv"
    #k_num_neighbors = 10

    # construct the argument and parse the arguments
    ap = argparse.ArgumentParser()
    ap.add_argument("-i", "--input_path", required = True, help = "path to *.ply model file")
    ap.add_argument("-o", "--output_path", required = True, help = "path to save *.csv file")
    ap.add_argument("-k", "--k_num_neighbors", required = True, type = int, help = "Number of nearest neighbors to use for feature calculation")
    args = vars(ap.parse_args())

    input_path = args["input_path"]
    output_path = args["output_path"]
    k_num_neighbors = args["k_num_neighbors"]
    
    feature_names = []
    for feature_domain in ['l','a','b']:
        for param in ["mean","std","entropy"]:
            feature_names.append(feature_domain + "_" + param)
    for feature_domain in ['curvature','anisotropy','linearity','planarity','sphericity']:
        for param in ["mean","std","entropy","ggd1","ggd2","aggd1","aggd2","aggd3","aggd4","gamma1","gamma2"]:
            feature_names.append(feature_domain + "_" + param)

    features_array = []
    try:
        result = get_feature_vector(input_path, k_num_neighbors)
    except:
        result = [None]*len(feature_names)
    features_array.append(result)

    df = pd.DataFrame(features_array,columns=feature_names, index=[input_path])
    df.sort_index(inplace=True)
    df.reset_index(inplace=True, names='path')
    df.to_csv(output_path, index=False, na_rep='NA')
```
