## Supplementary material for "Optimizing root phenotyping: Assessing the impact of camera calibration on 3D root reconstruction": filter_pointcloud.py

```
# Based on:
#
# Z. Zhang, W. Sun, X. Min, T. Wang, W. Lu and G. Zhai, 
# "No-Reference Quality Assessment for 3D Colored Point Cloud and Mesh Models," 
# in IEEE Transactions on Circuits and Systems for Video Technology, 
# vol. 32, no. 11, pp. 7618-7631, Nov. 2022, doi: 10.1109/TCSVT.2022.3186894.
# https://github.com/zzc-1998/NR-3DQA

"""
Statistical outlier removal

Author: Peter Pietrzyk
Date: December 20, 2023

"""

# import the necessary packages
import os
import sys
from pyntcloud import PyntCloud
import argparse

# Filter the ply model using Statistical Outlier Removal 
def model_filter_sor(model_file, output_path, k=10, z_max=3):
    
    if os.path.isfile(model_file):
        print("Filtering 3D point cloud model {}...".format(model_file))
    else:
        print("File does not exist")
        sys.exit()

    #load colored point cloud
    cloud = PyntCloud.from_file(model_file)

    # Statistical oulier removal
    kdtree = cloud.add_structure("kdtree")
    sor_filter = cloud.get_filter("SOR", kdtree_id=kdtree, k=k, z_max=z_max, and_apply=True)

    #write out point cloud file
    cloud.to_file(output_path)

    # check saved file
    if os.path.exists(output_path):
        print("Filtered 3D model was saved at {0}\n".format(output_path))
        return True
    else:
        print("Model filter failed!\n")
        return False

if __name__ == '__main__':
    
    # construct the argument and parse the arguments
    ap = argparse.ArgumentParser()
    ap.add_argument("-i", "--input_path", required = True, help = "path to *.ply model file")
    ap.add_argument("-o", "--output_path", required = True, help = "path to save filtered *.ply file")
    ap.add_argument("-k", "--k_num_neighbors", required = True, type = int, help = "Number of nearest neighbors to use")
    ap.add_argument("-z", "--z_max", required = True, type = float, help = "Maximum z score for SOR filter")
    args = vars(ap.parse_args())

    input_path = args["input_path"]
    output_path = args["output_path"]
    k_num_neighbors = args["k_num_neighbors"]
    z_max = args["z_max"]

    model_filter_sor(input_path, output_path, k=k_num_neighbors, z_max=z_max)
```
