## Supplementary material for "Optimizing root phenotyping: Assessing the impact of camera calibration on 3D root reconstruction": get_pointcloud_sizes.py

```
import argparse
import os 
import sys

import numpy as np
import pandas as pd
from pyntcloud import PyntCloud

"""
Script to get point cloud sizes of all files with specified file ending
Determines point cloud size of
    - *_clean.ply file
    - *_clean_filtered.ply file

Author: Peter Pietrzyk
Date: January 4, 2024
"""

def get_pc_size(model_file):
    if os.path.isfile(model_file):
        print("Reading 3D point cloud model {}...".format(model_file))
    else:
        print("File does not exist")
        sys.exit()

    cloud = PyntCloud.from_file(model_file)
    return len(cloud.points)

def find_files_with_extension(directory, file_extension):
    matching_files = []  # Initialize an empty list to store matching file paths

    # Walk through the directory and its subdirectories
    for dirpath, dirnames, filenames in os.walk(directory):
        for filename in filenames:
            if filename.endswith(file_extension):  # Check if the file ends with the specified extension
                file_path = os.path.join(dirpath, filename)  # Create the full file path
                file_path = file_path.replace("\\","/")
                matching_files.append(file_path)  # Add the file path to the list of matching files

    return matching_files  # Return the list of files with the specified extension

if __name__ == '__main__':

    """
    Example:
    python3 get_pointcloud_sizes.py -i "D:/UGA/Scanner Calibration/Paper/" -o "D:/UGA/Scanner Calibration/Analysis/pointcloud_sizes.csv -ext _clean.ply"
    """
    ap = argparse.ArgumentParser(description="Merge CSV files based on specified file extension.")
    ap.add_argument("-i", "--input_dir", required=True, help="Directory to search for CSV files")
    ap.add_argument("-o", "--output_path", required=True, help="Path to save merged CSV file")
    ap.add_argument("-ext", "--find_extension", required=True, help="File extension to search for")
    args = vars(ap.parse_args())

    input_dir = args["input_dir"]
    output_path = args["output_path"]
    find_extension = args["find_extension"]

    # Get column names and list of directories containing ply files
    file_list = find_files_with_extension(input_dir, find_extension)
    
    # Create empty dataframe
    df = pd.DataFrame(columns=['Pointcloud size'], index=file_list)

    # Add point cloud sizes to dataframe
    for file_path in df.index:                    
        pc_size = get_pc_size(file_path)
        df.at[file_path,'Pointcloud size'] = pc_size

        print(file_path, ' finished...')
    
    df = df.reset_index(names='path')
    df.to_csv(output_path, index=False, na_rep='NA')
```
