## Supplementary material for "Optimizing root phenotyping: Assessing the impact of camera calibration on 3D root reconstruction": nss_functions.py

```
# From:
#
# Z. Zhang, W. Sun, X. Min, T. Wang, W. Lu and G. Zhai, 
# "No-Reference Quality Assessment for 3D Colored Point Cloud and Mesh Models," 
# in IEEE Transactions on Circuits and Systems for Video Technology, 
# vol. 32, no. 11, pp. 7618-7631, Nov. 2022, doi: 10.1109/TCSVT.2022.3186894.
#
# Downloaded from https://github.com/zzc-1998/NR-3DQA


import numpy as np
import scipy
from scipy import stats
import pandas as pd
from scipy.stats import entropy
from scipy.special import gamma
from sklearn.preprocessing import scale,normalize
def my_scale(vec):
        vec = (vec-np.mean(vec))/np.std(vec, ddof=1)
        return vec

def get_color_nss_param(vec):
     """Estimate color NSS parameters.
        :param vec: The vector that we want to approximate its parameter.
        :type vec: np.ndarray
        :scale is the normalization function
        """
     return [estimate_basic_param(vec)]


def get_geometry_nss_param(vec):
     """Estimate geometry NSS parameters.
        :param vec: The vector that we want to approximate its parameter.
        :type vec: np.ndarray
        :scale is the normalization function
        """
     return [estimate_basic_param(vec),estimate_ggd_param(vec),estimate_aggd_param(my_scale(vec)),estimate_gamma_param(vec)]

def Entropy(labels):
    #probs = pd.Series(labels).value_counts() / len(labels)
    probs = pd.Series(labels).value_counts(bins = 2000) / len(labels)
    en = stats.entropy(probs)
    return en
    
    
def estimate_basic_param(vec):
        """Estimate basic parameter.
        :param vec: The vector that we want to approximate its parameter.
        :type vec: np.ndarray
        """
        result = [np.mean(vec),np.std(vec, ddof=1),Entropy(vec)]
        return result   
        
def estimate_ggd_param(vec):
        """Estimate GGD parameter.
        :param vec: The vector that we want to approximate its parameter.
        :type vec: np.ndarray
        """
        gam = np.arange(0.2, 10 + 0.001, 0.001)
        r_gam = (gamma(1.0 / gam) * gamma(3.0 / gam) / (gamma(2.0 / gam) ** 2))

        sigma_sq = np.mean(vec ** 2)
        sigma = np.sqrt(sigma_sq)
        E = np.mean(np.abs(vec))
        rho = sigma_sq / E ** 2

        differences = abs(rho - r_gam)
        array_position = np.argmin(differences)
        gamparam = gam[array_position]
        result = [gamparam, sigma]
        return result

def estimate_aggd_param(vec):
        """Estimate AGGD parameter.
        :param vec: The vector that we want to approximate its parameter.
        :type vec: np.ndarray
        """
        gam = np.arange(0.2, 10 + 0.001, 0.001)
        r_gam = ((gamma(2.0 / gam)) ** 2) / (
                    gamma(1.0 / gam) * gamma(3.0 / gam))

        left_std = np.sqrt(np.mean((vec[vec < 0]) ** 2))
        right_std = np.sqrt(np.mean((vec[vec > 0]) ** 2))
        gamma_hat = left_std / right_std
        rhat = (np.mean(np.abs(vec))) ** 2 / np.mean((vec) ** 2)
        rhat_norm = (rhat * (gamma_hat ** 3 + 1) * (gamma_hat + 1)) / (
                (gamma_hat ** 2 + 1) ** 2)

        differences = (r_gam - rhat_norm) ** 2
        array_position = np.argmin(differences)
        alpha = gam[array_position]
        const = np.sqrt(gamma(1 / alpha)) / np.sqrt(gamma(3 / alpha))
        mean_param = (right_std - left_std) * (
                        gamma(2 / alpha) / gamma(1 / alpha)) * const
        result = [alpha, mean_param,left_std, right_std]
        return result
       
            
def estimate_gamma_param(vec):
        """Estimate Gamma parameter.
        :param vec: The vector that we want to approximate its parameter.
        :type vec: np.ndarray
        """
        mean = np.mean(vec)
        std = np.std(vec)
        shape = (mean/std)**2
        scale = (std**2)/mean
        result = [shape,scale]
        return result
```
