## Supplementary material for "Optimizing root phenotyping: Assessing the impact of camera calibration on 3D root reconstruction": exp1_analyze_effect_of_calibration.py

```
import matplotlib.pyplot as plt
import numpy as np
import pandas as pd
from scipy import stats

"""
Performs statistical analysis of Experiment 1 and prints results.

Tests if calibration or SOR filter have effect on the 
five features (curvature, anisotropy, linearity, planarity, sphericity) 
or point cloud size

For results see: results_exp1_analyze_effect.txt 
"""

if __name__ == '__main__':

    # Paths
    data_path_pre_filtering = 'clean/200nn/table_clean_200nn.csv'
    data_path_post_filtering = 'clean_filtered/200nn/table_clean_filtered_200nn.csv'
    #table_save_path = 'results_exp1_analyze_effect.txt'
    
    # Read results
    # Unfiltered results
    df_unfiltered = pd.read_csv(data_path_pre_filtering)
    df_unfiltered['Filter'] = 'Unfiltered'

    # Unfiltered results
    df_filtered = pd.read_csv(data_path_post_filtering)
    df_filtered['Filter'] = 'Filtered'

    df_all = pd.concat([df_unfiltered, df_filtered], ignore_index=True)
    
    feature_list = ['curvature_mean','anisotropy_mean','linearity_mean','planarity_mean','sphericity_mean','Pointcloud size']
    label_list = ['mean (-)','mean (-)','mean (-)','mean (-)','mean (-)','number [#]']
    title_list = ['Curvature','Anisotropy','Linearity','Planarity','Sphericity','Pointcloud size']

    # Curvature
    sub_data = df_all.pivot(index='Root', columns=['Test', 'Filter'], values='curvature_mean')
    print('Curvature (mean)')
    print(sub_data.describe())
    print('Uncalibrated-Unfiltered vs Calibrated-Unfiltered')
    print(' - ', stats.wilcoxon(sub_data['E1#2: Uncalibrated']['Unfiltered'], sub_data['E1#1: Calibrated']['Unfiltered'])) # test calibrated-unfiltered effect
    print('Uncalibrated-Unfiltered vs Uncalibrated-Filtered')
    print(' - ', stats.wilcoxon(sub_data['E1#2: Uncalibrated']['Unfiltered'], sub_data['E1#2: Uncalibrated']['Filtered'])) # test unlibrated-filtered effect
    print('Uncalibrated-Unfiltered vs Calibrated-Filtered')
    print(' - ', stats.wilcoxon(sub_data['E1#2: Uncalibrated']['Unfiltered'], sub_data['E1#1: Calibrated']['Filtered']))   # test calibrated-filtered effect

    # Anisotropy
    sub_data = df_all.pivot(index='Root', columns=['Test', 'Filter'], values='anisotropy_mean')
    print('\nAnisotropy (mean)')
    print(sub_data.describe())
    print(' - ', 'Uncalibrated-Unfiltered vs Calibrated-Unfiltered')
    print(stats.wilcoxon(sub_data['E1#2: Uncalibrated']['Unfiltered'], sub_data['E1#1: Calibrated']['Unfiltered'])) # test calibrated-unfiltered effect
    print(' - ', 'Uncalibrated-Unfiltered vs Uncalibrated-Filtered')
    print(stats.wilcoxon(sub_data['E1#2: Uncalibrated']['Unfiltered'], sub_data['E1#2: Uncalibrated']['Filtered'])) # test unlibrated-filtered effect
    print(' - ', 'Uncalibrated-Unfiltered vs Calibrated-Filtered')
    print(stats.wilcoxon(sub_data['E1#2: Uncalibrated']['Unfiltered'], sub_data['E1#1: Calibrated']['Filtered']))   # test calibrated-filtered effect

    # Linearity
    sub_data = df_all.pivot(index='Root', columns=['Test', 'Filter'], values='linearity_mean')
    print('\nLinearity (mean)')
    print(sub_data.describe())
    print(' - ', 'Uncalibrated-Unfiltered vs Calibrated-Unfiltered')
    print(stats.wilcoxon(sub_data['E1#2: Uncalibrated']['Unfiltered'], sub_data['E1#1: Calibrated']['Unfiltered'])) # test calibrated-unfiltered effect
    print(' - ', 'Uncalibrated-Unfiltered vs Uncalibrated-Filtered')
    print(stats.wilcoxon(sub_data['E1#2: Uncalibrated']['Unfiltered'], sub_data['E1#2: Uncalibrated']['Filtered'])) # test unlibrated-filtered effect
    print(' - ', 'Uncalibrated-Unfiltered vs Calibrated-Filtered')
    print(stats.wilcoxon(sub_data['E1#2: Uncalibrated']['Unfiltered'], sub_data['E1#1: Calibrated']['Filtered']))   # test calibrated-filtered effect

    # Planarity
    sub_data = df_all.pivot(index='Root', columns=['Test', 'Filter'], values='planarity_mean')
    print('\nPlanarity (mean)')
    print(sub_data.describe())
    print(' - ', 'Uncalibrated-Unfiltered vs Calibrated-Unfiltered')
    print(stats.wilcoxon(sub_data['E1#2: Uncalibrated']['Unfiltered'], sub_data['E1#1: Calibrated']['Unfiltered'])) # test calibrated-unfiltered effect
    print(' - ', 'Uncalibrated-Unfiltered vs Uncalibrated-Filtered')
    print(stats.wilcoxon(sub_data['E1#2: Uncalibrated']['Unfiltered'], sub_data['E1#2: Uncalibrated']['Filtered'])) # test unlibrated-filtered effect
    print(' - ', 'Uncalibrated-Unfiltered vs Calibrated-Filtered')
    print(stats.wilcoxon(sub_data['E1#2: Uncalibrated']['Unfiltered'], sub_data['E1#1: Calibrated']['Filtered']))   # test calibrated-filtered effect

    # Sphericity
    sub_data = df_all.pivot(index='Root', columns=['Test', 'Filter'], values='sphericity_mean')
    print('\nSphericity (mean)')
    print(sub_data.describe())
    print(' - ', 'Uncalibrated-Unfiltered vs Calibrated-Unfiltered')
    print(stats.wilcoxon(sub_data['E1#2: Uncalibrated']['Unfiltered'], sub_data['E1#1: Calibrated']['Unfiltered'])) # test calibrated-unfiltered effect
    print(' - ', 'Uncalibrated-Unfiltered vs Uncalibrated-Filtered')
    print(stats.wilcoxon(sub_data['E1#2: Uncalibrated']['Unfiltered'], sub_data['E1#2: Uncalibrated']['Filtered'])) # test unlibrated-filtered effect
    print(' - ', 'Uncalibrated-Unfiltered vs Calibrated-Filtered')
    print(stats.wilcoxon(sub_data['E1#2: Uncalibrated']['Unfiltered'], sub_data['E1#1: Calibrated']['Filtered']))   # test calibrated-filtered effect

    # Point cloud size
    sub_data = df_all.pivot(index='Root', columns=['Test', 'Filter'], values='Pointcloud size')
    print('\nPoint cloud size')
    print(sub_data.describe())
    print(' - ', 'Uncalibrated-Unfiltered vs Calibrated-Unfiltered')
    print(stats.wilcoxon(sub_data['E1#2: Uncalibrated']['Unfiltered'], sub_data['E1#1: Calibrated']['Unfiltered'])) # test calibrated-unfiltered effect
    print(' - ', 'Uncalibrated-Unfiltered vs Uncalibrated-Filtered')
    print(stats.wilcoxon(sub_data['E1#2: Uncalibrated']['Unfiltered'], sub_data['E1#2: Uncalibrated']['Filtered'])) # test unlibrated-filtered effect
    print(' - ', 'Uncalibrated-Unfiltered vs Calibrated-Filtered')
    print(stats.wilcoxon(sub_data['E1#2: Uncalibrated']['Unfiltered'], sub_data['E1#1: Calibrated']['Filtered']))   # test calibrated-filtered effect
```
