## Supplementary material for "Optimizing root phenotyping: Assessing the impact of camera calibration on 3D root reconstruction": exp1_export_pc_with_features.py

```
import argparse
from pyntcloud import PyntCloud

"""
Exports a point cloud with feature as scalar values for each point.
"""
def get_feature_vector(objpath, outputpath, k):  
    #load colored point cloud
    print("Begin loading point cloud")
    cloud = PyntCloud.from_file(objpath)
    
    #begin geometry projection
    print("Begin geometry feature extraction.")
    k_neighbors = cloud.get_neighbors(k=k)
    ev = cloud.add_scalar_field("eigen_values", k_neighbors=k_neighbors)
    cloud.add_scalar_field("curvature", ev=ev)
    cloud.add_scalar_field("anisotropy",ev=ev)
    cloud.add_scalar_field("linearity",ev=ev)
    cloud.add_scalar_field("planarity",ev=ev)
    cloud.add_scalar_field("sphericity",ev=ev)
    
    # Set min and max values for better scaling
    kplusone = k+1
    cloud.points['curvature({})'.format(kplusone)][0] = 0
    cloud.points['curvature({})'.format(kplusone)][1] = 1
    cloud.points['anisotropy({})'.format(kplusone)][0] = 0
    cloud.points['anisotropy({})'.format(kplusone)][1] = 1
    cloud.points['linearity({})'.format(kplusone)][0] = 0
    cloud.points['linearity({})'.format(kplusone)][1] = 1
    cloud.points['planarity({})'.format(kplusone)][0] = 0
    cloud.points['planarity({})'.format(kplusone)][1] = 1
    cloud.points['sphericity({})'.format(kplusone)][0] = 0
    cloud.points['sphericity({})'.format(kplusone)][1] = 1
    
    """
    curvature = cloud.points['curvature({})'.format(kplusone)].to_numpy()
    anisotropy = cloud.points['anisotropy({})'.format(kplusone)].to_numpy()
    linearity = cloud.points['linearity({})'.format(kplusone)].to_numpy()
    planarity = cloud.points['planarity({})'.format(kplusone)].to_numpy()
    sphericity = cloud.points['sphericity({})'.format(kplusone)].to_numpy()
    """
    cloud.to_file(outputpath)    

if __name__ == '__main__':

    # construct the argument and parse the arguments
    ap = argparse.ArgumentParser()
    ap.add_argument("-i", "--input_path", required = True, help = "path of reference *.ply model file")
    ap.add_argument("-o", "--output_path", required = True, help = "path for output .ply file with features")
    ap.add_argument("-k", "--k_num_neighbors", required = True, type = int, help = "Number of nearest neighbors to use for feature calculation")

    args = vars(ap.parse_args())

    input_path = args["input_path"]
    output_path = args["output_path"]
    k_num_neighbors = args["k_num_neighbors"]
    
    get_feature_vector(input_path, output_path, k_num_neighbors)
```
