## Supplementary material for "Optimizing root phenotyping: Assessing the impact of camera calibration on 3D root reconstruction": exp1_plot_effect_of_calibration.py

```
import os
import argparse
import math

import matplotlib.pyplot as plt
import numpy as np
import pandas as pd
from scipy import stats

"""
Creates a plot showing how features change in each root when scanner is calibrated.
"""

def plot_paired(vals1, vals2, label, ax_id):

    # plotting the points
    ax_id.scatter(np.zeros(len(vals1)), vals1, c='k')
    ax_id.scatter(np.ones(len(vals2)), vals2, c='k')

    # plotting the lines
    for i in range(len(vals1)):
        ax_id.plot( [0,1], [vals1[i], vals2[i]], c='k')
        ax_id.set_xlim([-0.15,1.15])
    ax_id.set_xticks([0,1], ['Uncalib.', 'Calib.'])
    ax_id.tick_params(which="minor", axis="y", direction="inout")
    

if __name__ == '__main__':

    # Paths
    data_path = 'clean/200nn/table_clean_200nn.csv'
    save_path = 'clean/calibration_clean_200nn.png'
    
    # Read results
    df = pd.read_csv(data_path)
    df = df.loc[(df['Test'] == 'E1#1: Calibrated') | (df['Test'] == 'E1#2: Uncalibrated')]
    feature_list = ['curvature_mean','anisotropy_mean','linearity_mean','planarity_mean','sphericity_mean','Pointcloud size']
    label_list = ['mean (-)','mean (-)','mean (-)','mean (-)','mean (-)','number [#]']
    title_list = ['Curvature','Anisotropy','Linearity','Planarity','Sphericity','Pointcloud size']

    fig, ax = plt.subplots(1, len(feature_list), figsize=(14, 3))

    # Curvature
    sub_data = df.pivot(index='Root', columns='Test', values='curvature_mean')
    plot_paired(sub_data['E1#2: Uncalibrated'], sub_data['E1#1: Calibrated'], 'mean (-)', ax[0])
    ax[0].set_ylim([0.143, 0.191])
    ax[0].set_yticks(np.arange(0.15, 0.195, 0.01))
    ax[0].set_ylabel('mean (-)')
    ax[0].set_title('Curvature (k=200)')
    print('Curvature (mean): E1#1: Calibrated vs E1#2: Uncalibrated')
    print(stats.wilcoxon(sub_data['E1#2: Uncalibrated'], sub_data['E1#1: Calibrated']))
    print(stats.ttest_rel(sub_data['E1#2: Uncalibrated'], sub_data['E1#1: Calibrated']))

    # Anisotropy
    sub_data = df.pivot(index='Root', columns='Test', values='anisotropy_mean')
    plot_paired(sub_data['E1#2: Uncalibrated'], sub_data['E1#1: Calibrated'], 'mean (-)', ax[1])
    ax[1].set_ylim([0.575, 0.705])
    ax[1].set_yticks(np.arange(0.58, 0.705, 0.02))
    ax[1].set_ylabel('mean (-)')
    ax[1].set_title('Anisotropy (k=200)')
    print('\nAnisotropy (mean): E1#1: Calibrated vs E1#2: Uncalibrated')
    print(stats.wilcoxon(sub_data['E1#2: Uncalibrated'], sub_data['E1#1: Calibrated']))
    print(stats.ttest_rel(sub_data['E1#2: Uncalibrated'], sub_data['E1#1: Calibrated']))

    # Linearity
    sub_data = df.pivot(index='Root', columns='Test', values='linearity_mean')
    plot_paired(sub_data['E1#2: Uncalibrated'], sub_data['E1#1: Calibrated'], 'mean (-)', ax[2])
    ax[2].set_ylim([0.235, 0.305])
    ax[2].set_yticks(np.arange(0.24, 0.305, 0.01))
    ax[2].set_ylabel('mean (-)')
    ax[2].set_title('Linearity (k=200)')
    print('\nLinearity (mean): E1#1: Calibrated vs E1#2: Uncalibrated')
    print(stats.wilcoxon(sub_data['E1#2: Uncalibrated'], sub_data['E1#1: Calibrated']))
    print(stats.ttest_rel(sub_data['E1#2: Uncalibrated'], sub_data['E1#1: Calibrated']))

    # Planarity
    sub_data = df.pivot(index='Root', columns='Test', values='planarity_mean')
    plot_paired(sub_data['E1#2: Uncalibrated'], sub_data['E1#1: Calibrated'], 'mean (-)', ax[3])
    ax[3].set_ylim([0.335, 0.425])
    ax[3].set_yticks(np.arange(0.34, 0.425, 0.01))
    ax[3].set_ylabel('mean (-)')
    ax[3].set_title('Planarity (k=200)')
    print('\nPlanarity (mean): E1#1: Calibrated vs E1#2: Uncalibrated')
    print(stats.wilcoxon(sub_data['E1#2: Uncalibrated'], sub_data['E1#1: Calibrated']))
    print(stats.ttest_rel(sub_data['E1#2: Uncalibrated'], sub_data['E1#1: Calibrated']))

    # Sphericity
    sub_data = df.pivot(index='Root', columns='Test', values='sphericity_mean')
    plot_paired(sub_data['E1#2: Uncalibrated'], sub_data['E1#1: Calibrated'], 'mean (-)', ax[4])
    ax[4].set_ylim([0.295, 0.425])
    ax[4].set_yticks(np.arange(0.30, 0.425, 0.02))
    ax[4].set_ylabel('mean (-)')
    ax[4].set_title('Sphericity (k=200)')
    print('\nSphericity (mean): E1#1: Calibrated vs E1#2: Uncalibrated')
    print(stats.wilcoxon(sub_data['E1#2: Uncalibrated'], sub_data['E1#1: Calibrated']))
    print(stats.ttest_rel(sub_data['E1#2: Uncalibrated'], sub_data['E1#1: Calibrated']))

    # Point cloud size
    sub_data = df.pivot(index='Root', columns='Test', values='Pointcloud size')
    plot_paired(sub_data['E1#2: Uncalibrated']/1e6, sub_data['E1#1: Calibrated']/1e6, 'mean (-)', ax[5])
    ax[5].set_ylim([2.8, 8.1])
    ax[5].set_yticks(np.arange(3, 8.1, 1))
    ax[5].set_ylabel('number (in millions)')
    ax[5].set_title('Point cloud size')
    print('\nPoint cloud size: E1#1: Calibrated vs E1#2: Uncalibrated')
    print(stats.wilcoxon(sub_data['E1#2: Uncalibrated'], sub_data['E1#1: Calibrated']))
    print(stats.ttest_rel(sub_data['E1#2: Uncalibrated'], sub_data['E1#1: Calibrated']))

    #plt.show()
    fig.tight_layout()
    plt.savefig(save_path, dpi=600)
```
