## Supplementary material for "Optimizing root phenotyping: Assessing the impact of camera calibration on 3D root reconstruction": exp1_plot_effect_of_k.py

```
import os
import argparse
import math

import matplotlib.pyplot as plt
import numpy as np
import pandas as pd
import seaborn as sns
    
"""
Plots the effect of number of neighbors 'k' in each root model for values from k=10 to k=400.
"""

if __name__ == '__main__':

    # Load all tables for k=10 to 200
    data_path_10 = 'clean/10nn/table_clean_10nn.csv'
    df_10 = pd.read_csv(data_path_10)
    df_10 = df_10.loc[(df_10['Test'] == 'E1#1: Calibrated')]
    #df_10['k'] = 10

    data_path_25 = 'clean/25nn/table_clean_25nn.csv'
    df_25 = pd.read_csv(data_path_25)
    df_25 = df_25.loc[df_25['Test'] == 'E1#1: Calibrated']
    #df_25['k'] = 25

    data_path_50 = 'clean/50nn/table_clean_50nn.csv'
    df_50 = pd.read_csv(data_path_50)
    df_50 = df_50.loc[df_50['Test'] == 'E1#1: Calibrated']
    #df_50['k'] = 50

    data_path_100 = 'clean/100nn/table_clean_100nn.csv'
    df_100 = pd.read_csv(data_path_100)
    df_100 = df_100.loc[df_100['Test'] == 'E1#1: Calibrated']
    #df_100['k'] = 100

    data_path_200 = 'clean/200nn/table_clean_200nn.csv'
    df_200 = pd.read_csv(data_path_200)
    df_200 = df_200.loc[df_200['Test'] == 'E1#1: Calibrated']
    #df_200['k'] = 200

    data_path_300 = 'clean/300nn/table_clean_300nn.csv'
    df_300 = pd.read_csv(data_path_300)
    df_300 = df_300.loc[df_300['Test'] == 'E1#1: Calibrated']
    #df_300['k'] = 300

    data_path_400 = 'clean/400nn/table_clean_400nn.csv'
    df_400 = pd.read_csv(data_path_400)
    df_400 = df_400.loc[df_400['Test'] == 'E1#1: Calibrated']
    #df_400['k'] = 400

    save_path = 'clean/effect_of_k_clean.png'
    custom_order = ['root_1', 'root_2', 'root_3', 'root_4', 'root_5', 'root_6', 'root_7', 'root_8', 'root_9', 'root_10', 'root_11', 'root_12']

    feature_list = ['curvature_mean','anisotropy_mean','linearity_mean','planarity_mean','sphericity_mean']
    df_all = pd.concat([df_10, df_25, df_50, df_100, df_200, df_300, df_400], ignore_index=True)
    df_all = df_all.melt(id_vars=['Root', 'k'], value_vars=feature_list)
    sns.relplot(data=df_all, x='k', y='value', hue='Root', hue_order=custom_order, col='variable', kind="line", height=3, facet_kws={'sharey': False, 'sharex': True})
    #plt.show()
    plt.savefig(save_path, dpi=600)
```
