## Supplementary material for "Optimizing root phenotyping: Assessing the impact of camera calibration on 3D root reconstruction": exp1_plot_kde.py

```
"""
Created by: Peter Pietrzyk


December 21, 2023

Creates kdf plots
"""

import os 
import argparse
import matplotlib.pyplot as plt
import matplotlib.ticker as ticker
import seaborn as sns
import numpy as np
import pandas as pd
from skimage import color
from pyntcloud import PyntCloud

from nss_functions import * 

"""
Plots histograms with distribution of all five feature for selected root models.
Two root models can be selected in order to be compared.
"""

def get_feature_vector(objpath, k):  
    #load colored point cloud
    print("Begin loading point cloud")
    cloud = PyntCloud.from_file(objpath)
    
    #begin geometry projection
    print("Begin geometry feature extraction.")
    k_neighbors = cloud.get_neighbors(k=k)
    ev = cloud.add_scalar_field("eigen_values", k_neighbors=k_neighbors)
    cloud.add_scalar_field("curvature", ev=ev)
    cloud.add_scalar_field("anisotropy",ev=ev)
    cloud.add_scalar_field("linearity",ev=ev)
    cloud.add_scalar_field("planarity",ev=ev)
    cloud.add_scalar_field("sphericity",ev=ev)
    kplusone = k+1
    curvature = cloud.points['curvature({})'.format(kplusone)].to_numpy()
    anisotropy = cloud.points['anisotropy({})'.format(kplusone)].to_numpy()
    linearity = cloud.points['linearity({})'.format(kplusone)].to_numpy()
    planarity = cloud.points['planarity({})'.format(kplusone)].to_numpy()
    sphericity = cloud.points['sphericity({})'.format(kplusone)].to_numpy()

    output = [curvature,anisotropy,linearity,planarity,sphericity]
    return output

def subsample_feature_array(features_array, num_samples):
    """
    Subsample the features_array by randomly selecting a specified number of samples.
    The same set of samples is selected across all sub-arrays.
    
    Parameters:
        features_array (list of lists): List containing feature arrays.
        num_samples (int): Number of samples to subsample from the feature arrays.
    
    Returns:
        subsampled_features_array (list of lists): Subsampled feature arrays.
    """
    # Ensure the number of samples is not greater than the number of available samples
    num_samples = min(num_samples, len(features_array[0]))
    
    # Randomly select indices for subsampling
    indices = np.random.choice(len(features_array[0]), size=num_samples, replace=False)
    
    # Apply the same set of indices to all feature arrays
    subsampled_features_array = [feature_array[indices] for feature_array in features_array]
    
    return subsampled_features_array


def plot_pair(ref_pc, compare_pc, k, save_path, num_samples=None):
    """
    Function for plotting pairwise comparisons between features of two point clouds.
    
    Parameters:
        ref_pc (list of lists): List containing feature arrays from the reference point cloud.
        compare_pc (list of lists): List containing feature arrays from the comparison point cloud.
        k (int): Number of nearest neighbors.
        save_path (str): Path where the combined plot will be saved.
        num_samples (int, optional): Number of samples to subsample from the feature arrays. Default is None.
    """
    # If num_samples is provided, subsample the feature arrays
    if num_samples is not None:
        ref_pc = subsample_feature_array(ref_pc, num_samples)
        compare_pc = subsample_feature_array(compare_pc, num_samples)

    # Feature labels
    x_labels = ['Curvature (k={})'.format(str(k)), 'Anisotropy (k={})'.format(str(k)), 'Linearity (k={})'.format(str(k)), 'Planarity (k={})'.format(str(k)), 'Sphericity (k={})'.format(str(k))]

    # Create subplots for each feature
    fig, axes = plt.subplots(1, len(ref_pc), figsize=(8, 2))

    for i, (ref_feature, comp_feature) in enumerate(zip(ref_pc, compare_pc)):
        ax = axes[i]
        
        # Plot KDE for reference and comparison features
        #sns.kdeplot(ref_feature, ax=ax, label='Calibrated')
        #sns.kdeplot(comp_feature, ax=ax, label='Uncalibrated')

        bin_width = 0.005  # Adjust this value as needed
        bins = np.arange(0, 1 + bin_width, bin_width)

        # Plot histograms for reference and comparison features
        ax.hist(ref_feature, bins=bins, zorder=2, alpha=0.5, label='Calibrated')
        ax.hist(comp_feature, bins=bins, zorder=1, alpha=0.5, label='Uncalibrated')

        # Plot vertical lines at the mean of each distribution
        ax.axvline(np.mean(ref_feature), color='blue', linestyle='dashed', linewidth=0.75, label='Mean (Calib.)')
        ax.axvline(np.mean(comp_feature), color='orange', linestyle='dashed', linewidth=0.75, label='Mean (Uncalib.)')

        # Set x ticks
        if i == 0:
            ax.set_xlim([0, 0.33])
            ax.set_xticks([0, 0.33])
        else:
            ax.set_xlim([0, 1])

        # Set y label for the left plot only
        if i == 0:
            ax.set_ylabel('Density', fontsize=8)
        else:
            ax.set_ylabel('')

        # Set x label
        ax.set_xlabel(x_labels[i], fontsize=8)

        # Decrease font size
        ax.tick_params(axis='both', which='major', labelsize=6)
        
        ax.yaxis.set_major_formatter(ticker.ScalarFormatter())
        ax.yaxis.major.formatter.set_powerlimits((0, 0))
        ax.yaxis.offsetText.set_fontsize(6)

    # Add legend to the last subplot
    ax.legend(loc='center left', bbox_to_anchor=(1, 0.5), fontsize=8)

    # Adjust layout to prevent overlap
    plt.tight_layout()

    # Save the combined plot
    plt.savefig(save_path, dpi=600)
    plt.close()


if __name__ == '__main__':
    """
    reference_path = "E1_1_Calibrated/root_1_good/root_1_good_clean.ply"
    compare_path = "E1_1_Calibrated/root_1_good/root_1_bad/root_1_bad_clean.ply"
    output_path = "comparison_kde_plot.png"
    k_num_neighbors = 25

    Example
    python exp1_plot_kde.py -ref "E1_1_Calibrated/root_1_good/root_1_good_clean.ply" -com "E1_1_Calibrated/root_1_good/root_1_bad/root_1_bad_clean.ply" -o comparison_kde_plot.png -k 25
    """
    
    # Argument parsing
    ap = argparse.ArgumentParser()
    ap.add_argument("-ref", "--reference_path", required=True, help="path of reference *.ply model file")
    ap.add_argument("-com", "--compare_path", required=True, help="path of compared *.ply model file")
    ap.add_argument("-o", "--output_path", required=True, help="Path to save *.png image")
    ap.add_argument("-k", "--k_num_neighbors", required=True, type=int, help="Number of nearest neighbors to use for feature calculation")
    args = vars(ap.parse_args())

    # Extract arguments
    reference_path = args["reference_path"]
    compare_path = args["compare_path"]
    output_path = args["output_path"]
    k_num_neighbors = args["k_num_neighbors"]
    

    # Get feature vectors from point clouds
    features_array_ref = get_feature_vector(reference_path, k=k_num_neighbors)
    features_array_comp = get_feature_vector(compare_path, k=k_num_neighbors)

    # Combine features into a single plot
    plot_pair(features_array_ref, features_array_comp, k_num_neighbors, output_path)
```
