## Supplementary material for "Optimizing root phenotyping: Assessing the impact of camera calibration on 3D root reconstruction": exp2_plot_heatmaps.py

```
"""
Created by: Peter Pietrzyk


December 8, 2023

Reads csv file with results and plots a heatmap that show effect a change in camera settings on features.
"""
import numpy as np
import pandas as pd
import os 
import matplotlib.pyplot as plt
import seaborn as sns

# Custom function to format values
def format_mean_std(mean, std_dev):
    return '{:.2f} ({:.2f})'.format(mean, std_dev)


# INPUT/OUTPUT
# Path for reading data table
data_path = 'clean/200nn/table_clean_200nn.csv'

# Extension for figure filenames
figure_name = 'heatmap_experiment2_200nn.png'

# Directory to save plots
dir_save_plots = 'clean/'

# Read data
df_merged = pd.read_csv(data_path)

# PREPARE DATA
# List all features 
feature_names = ['curvature_mean','anisotropy_mean','linearity_mean','planarity_mean','sphericity_mean','Pointcloud size']
rename_features = {'curvature_mean':'Curvature','anisotropy_mean':'Anisotropy','linearity_mean':'Linearity','planarity_mean':'Planarity','sphericity_mean':'Sphericity','Pointcloud size':'Point cloud size'}
# Drop unneccesary columns
df_merged = df_merged[['Root','Test']+feature_names].set_index(['Root', 'Test'])
df_merged.rename(columns=rename_features, inplace=True)

test_order = ['E2#01: Calibrated',
                   'E2#02: Jpg 100',
                   'E2#03: 1/125sec',
                   'E2#04: 1/60sec',
                   'E2#05: 1/15sec',
                   'E2#06: 1/8sec',
                   'E2#07:1/125sec, +6dB',
                   'E2#08: 1/500sec, +12dB',
                   'E2#09: f/2, 1/125sec',
                   'E2#10: f/8, 1/8sec',
                   'E2#11: f/8, 1/15sec, +3dB',
                   'E2#12: f/8, 1/30sec, +6dB',
                   'E2#13: +/-10cm (#1)',
                   'E2#14: +/-10cm (#2)',
                   'E2#15: +/-10cm, f/2, 1/125sec (#1)',
                   'E2#16: +/-10cm, f/2, 1/125sec (#2)',
                   'E2#17: +/-10cm, f/8, 1/15sec, +3dB (#1)',
                   'E2#18: +/-10cm, f/8, 1/15sec, +3dB (#2)',
                   'E2#19: +/-17cm',
                   'E2#20: +/-17cm, f/2, 1/125sec',
                   'E2#21: +/-17cm, f/8, 1/15sec, +3dB',
                   'E2#22: Distance +10cm',
                   'E2#23: Camera orientation',
                   'E2#24: Root position']


# Normalize 
df_reference = df_merged[df_merged.index.get_level_values('Test') == 'E2#01: Calibrated'] 
df_reference = df_reference.droplevel(level='Test')
df_merged = df_merged.sub(df_reference, level = 'Root').div(df_reference, level = 'Root') * 100

# Get mean
df_merged_means = df_merged.groupby('Test').mean().reindex(test_order)
#df_merged_means = df_merged_means.reset_index().set_index('Test').reindex(test_order)

# Get standard deviation
df_merged_std = df_merged.groupby('Test').std().reindex(test_order)
#df_merged_std = df_merged_std.reset_index().set_index('Test').reindex(test_order)

df_merged_combined = df_merged_means.applymap('{:,.1f}'.format).add('%\n (').add(df_merged_std.applymap('{:,.1f}'.format)).add(')')

save_plot_path = os.path.join(dir_save_plots, figure_name)

fig = plt.figure(figsize = (8,8))
sns.heatmap(df_merged_means, vmin=-20, vmax=+20, cmap='vlag', square=False, annot=df_merged_combined, fmt="", annot_kws = {'fontsize':8}, linecolor='k', linewidth=.5)

fig.tight_layout()
plt.savefig(save_plot_path,dpi=600)
plt.close(fig)
```
